# Enhancer binding kinetics explain transcription factor hub formation

**DOI:** 10.1101/2025.04.07.647578

**Authors:** Samantha Fallacaro, Manya Kapoor, Lillian Encarnation, Apratim Mukherjee, Meghan A. Turner, Hernan G. Garcia, Mustafa Mir

## Abstract

Transcription factors (TFs) form dynamic, high-concentration clusters, condensates, or hubs, proposed to increase TF binding frequency at target enhancers. However, how enhancer sequence shapes hub properties remains unclear. We developed a live-imaging based framework to quantify the spatiotemporal relationship between TF hubs and actively transcribed genes in live Drosophila embryos. Examining hubs formed by the TF Dorsal across enhancers with defined binding-site composition, we find that hub enrichment and persistence scale with the number of Dorsal binding motifs. However, these hub properties do not predict transcriptional bursts for a given enhancer. Combining quantitative imaging with computational modeling, we show that Dorsal hub formation can be explained by TF–DNA binding kinetics alone. These findings support a model in which TF hubs emerge from enhancer-encoded TF–DNA interactions rather than higher-order regulatory assemblies.

## Introduction

The spatiotemporal regulation of gene expression during embryonic development depends on the ability of sequence-specific transcription factors to find and bind their target sites within the crowded nuclear environment. Live-imaging and single-molecule tracking experiments have shown that transcription factors (TFs) bind their genomic targets transiently, with residence times on the order of tens of seconds (*1, 2*). The local accumulation of TFs around targets, through the formation of hubs or condensates, has been proposed as a mechanism to increase binding frequency and maintain target occupancy despite short residence times (*1–7*). These high-local concentration clusters are further thought to catalyze the recruitment of co-factors and transcriptional machinery to amplify gene activation (*8–22*). Here, we sought to address the central question of this model: whether hubs drive TF target occupancy by increasing their binding frequency or whether they are an emergent feature of underlying binding kinetics.

Clusters of transcription factors are referred to as condensates when they exhibit non-stochiometric accumulations and their biophysical properties are consistent with formation via phase separation mechanisms (*23–25*). Under the phase separation model, transcriptional condensate formation depends on the nuclear concentration and sequence properties of condensate constituents. Although substantial work has focused on how intrinsic protein features and concentration thresholds govern condensate formation (*7*), far less attention has been directed toward how genomic context may tune hub properties at individual loci in an *in vivo* context.

Hub localization to genomic sites is dynamically influenced by a combination of targeting through a TF’s protein-protein and protein-DNA interactions (*11, 16, 21, 26–30*). A TF’s locus-specific network of interaction partners also plays a role in generating compositionally and functionally distinct hubs (*27, 30, 31*). The nuclear concentration of a TF and the number of TF binding sites at a locus has a direct impact on TF occupancy and transcriptional output (*11, 32*). Recent work has suggested a spatio-temporal relationship where the proximity of a hub to its regulatory elements can increase transcriptional bursting (*16, 33–35*). If hubs differentially influence TF occupancy at genes, their properties should be highly gene-dependent, rather than globally dictated by transcription factor identity and nuclear concentration. Based on these observations we hypothesized that the biophysical properties attributed to hubs, such as their size and stability, are not emergent properties of higher-order phenomena such as phase-separation. We reasoned that hub properties instead reflect the molecular scale binding and diffusion kinetics of hub constituents in an enhancer-specific manner.

To investigate whether hubs drive or reflect binding kinetics, we leverage the *Drosophila* dorsoventral patterning system. In this system, the transcription factor Dorsal forms a nuclear concentration gradient and activates distinct sets of genes in a spatially-dependent manner along the embryo’s dorsoventral axis (*36, 37*). Due to its morphogenic nature, Dorsal-regulated gene expression provides a natural framework to determine whether hubs function uniformly or differentially at target genes that are expressed at different Dorsal concentration levels. Previous work in the *Drosophila* embryo has shown that TF hubs containing Bicoid, Zelda, or Dorsal are recruited to specific genomic loci even at low concentrations, and suggests that morphogen hubs play an active role in fine-tuning transcriptional responses to spatial gradients (*8, 9, 21, 32, 38, 39*). Dorsal clusters have recently been proposed to form in a concentration-threshold-dependent manner via liquid-liquid phase separation mechanisms (*40*). However, how these clusters associate with target sites and if their properties are tuned in a site-specific manner has not previously been explored.

Here, we use high-resolution lattice light-sheet imaging to analyze how Dorsal hubs form at and interact with transcriptionally active loci. To enable these insights, we developed a live imaging-based framework that quantifies the spatiotemporal relationship between TF hubs and active transcription sites *in vivo*. Using this approach, we measured Dorsal hub properties including local Dorsal enrichment and the persistence of hubs at enhancers of *sog* and *snail*, and at synthetic enhancers with varying numbers of Dorsal binding sites. We further used our live-imaging data, along with single-molecule tracking experiments, to parameterize a molecular kinetics model that recapitulates Dorsal hub formation at the enhancers we investigated. Together, our results suggest that TF clustering, or hub formation, primarily reflects enhancer-dependent binding kinetics rather than a distinct regulatory layer that increases binding frequency.

## Results

### Dorsal hub persistence and density scale with nuclear concentration

To investigate the biophysical properties of Dorsal hubs, we performed lattice light-sheet imaging on blastoderm stage *Drosophila* embryos expressing endogenously-tagged Dorsal-mNeonGreen in the 13th and 14th syncytial nuclear cycles (nc13 and nc14) (*41*). Dorsal forms a nuclear concentration gradient and is found at high concentrations in ventral nuclei, moderate to low concentrations in lateral nuclei, and is largely excluded from dorsal nuclei (Fig. 1A, Movie S1). Dorsal’s nuclear concentration gradient is not persistent and is re-established after each nuclear division. At the onset of mitosis, the cytoplasmic-nuclear concentration ratio flattens, and after nuclear membrane reformation the gradient is re-established through a balance of nuclear import and export during interphase (*37*). We observed that Dorsal hubs are visible as soon as Dorsal accumulates in the nucleus after mitosis and remain visible throughout interphase at nuclei in both ventral and lateral regions (Fig. 1B-C). Further, hubs form even as the nucleoplasmic Dorsal concentration is diminished in the dorsal-lateral region (Fig. S1A,B, Movie S2), which demonstrates that hub formation does not require high nuclear protein levels.

**Figure 1:**
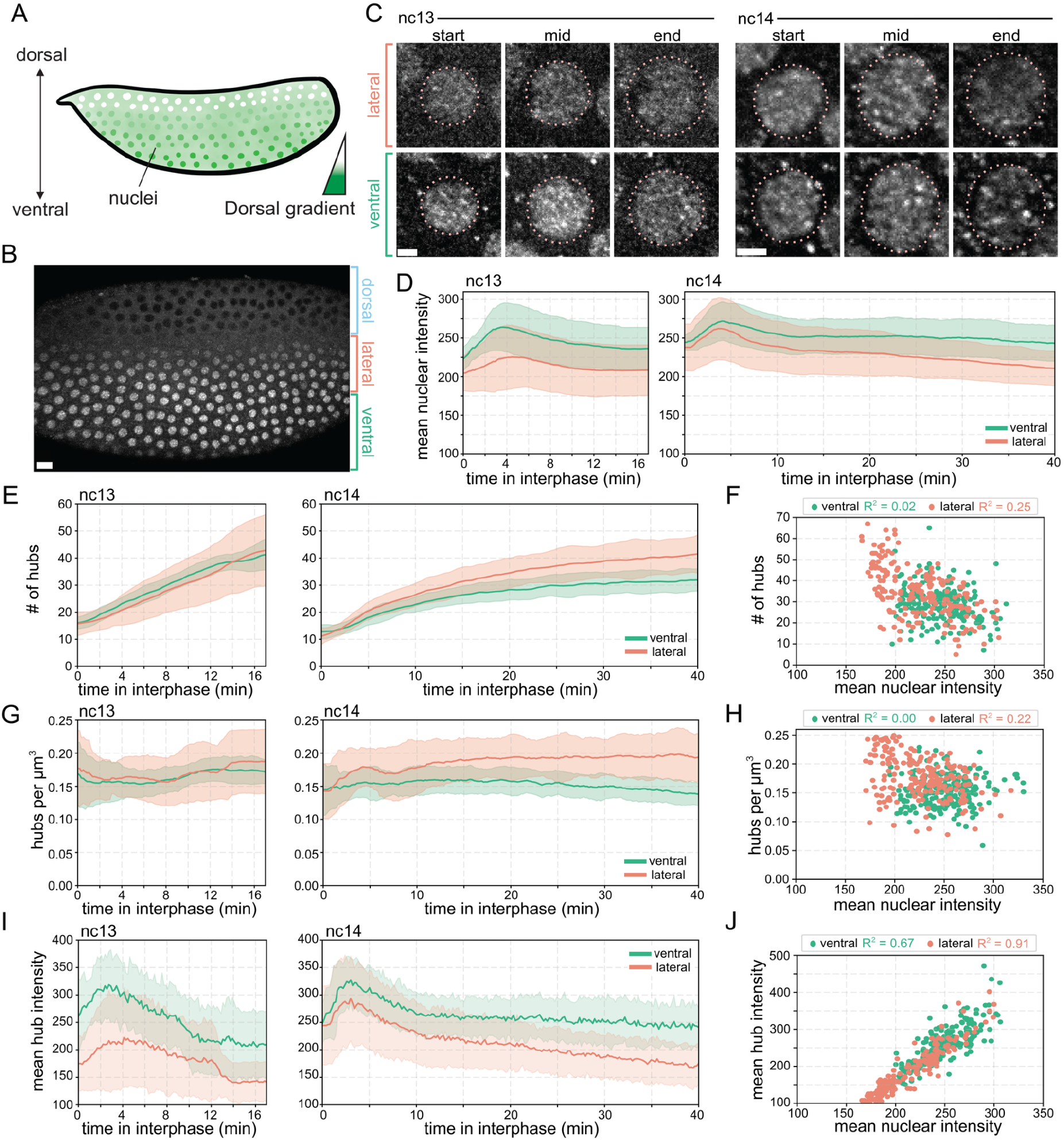
Dorsal hub densities are constant in interphase, but hub enrichment depends on Dorsal nuclear concentration. **(A)** Schematic of the Dorsal gradient in *Drosophila* embryos. Dorsal translocates into ventral surface nuclei at higher rates creating a nuclear concentration gradient along the dorsoventral axis. **(B)** Maximum intensity projection image showing nuclear gradient of Dorsal-mNeonGreen in a *Drosophila* embryo in nc13. Scale bar is 15 microns. **(C)** Representative single z-slice images of Dorsal-mNeonGreen in nuclei at the lateral (top) and ventral (bottom) embryonic surfaces at the start, middle, and end of nc13 (left) and nc14 (right). Scale bars are 2 microns. **(D)** Average nuclear mean intensity in ventral (green) or lateral (orange) nuclei. Shading shows standard deviation between embryo replicates. N = 5 nuclei per embryo, 3 embryo replicates for each axis position. **(E)** Average number of hubs per nucleus in ventral (green) or lateral (orange) nuclei. Shading shows standard deviation between embryo replicates. **(F)** Scatter plot of number of hubs vs. mean nuclear intensities. R^2^ value is Pearson’s correlation (N = 5814 ventral hubs, 6928 lateral hubs from N = 3 embryos per axis position). **(G)** Average hub density (number of hubs per cubic micron) in ventral (green) or lateral (orange) nuclei. Shading shows standard deviation between embryo replicates. **(H)** Scatter plot of hub densities vs. mean nuclear intensities. R^2^ value is Pearson’s correlation. **(I)** Mean intensities of Dorsal hubs in ventral (green) or lateral (orange) nuclei. Solid line shows average intensity and shading shows standard deviation between embryo replicates. **(J)** Scatter plot showing hub mean intensity and mean nuclear intensity. R^2^ is Pearson’s correlation.

To understand how nuclear concentrations influence hub formation, we compared ventral nuclei along the presumptive ventral furrow with nuclei at the lateral surface (Fig. 1B). Here, we define lateral nuclei as the region of the embryo where Dorsal nuclear concentration diminishes to be equivalent to the cytoplasmic concentration (Fig. S1A,B). Nuclear Dorsal intensity in both ventral and lateral nuclei rises for the first 4.5 minutes of interphase and then falls for the remainder of interphase (Fig. 1D). To compare Dorsal hubs in these nuclei, we developed a custom analysis pipeline to detect hubs and quantify their intensities, spatial distributions, and persistence (Fig. S1C, Movie S3). While the total number of detected hubs per nucleus increases during interphase (Fig. 1E-F), this increase is correlated with the expansion of nuclear volume in this time period (Fig. S1D-E). Thus, despite the dynamic change in nuclear concentration during interphase, the density of Dorsal hubs (number of hubs per unit volume) remains relatively constant during the interphases of nc13 and nc14 (Figs. 1G-H). We suspect that the apparent increase in hub number is due to tightly spaced accumulations at binding sites becoming resolvable as the nuclear volume expands rather than *de novo* formation of additional hubs.

Although the density of hubs is uncoupled with nuclear concentration, the average hub intensity is strongly correlated with nuclear Dorsal concentration (Figs. 1I-J, S1H). To further assess hub variability using a segmentation-independent metric, we calculated the coefficient of variation (CV) of nuclear Dorsal intensity (Fig. S1F-G). The CV exhibits an intermediate correlation with nuclear concentration likely since the CV reflects intensity dispersion and not just morphological differences. Based on these data we hypothesized that hubs form rapidly at dorsal target sites at the start of interphase, and then increasing amounts of Dorsal interact with these sites as nuclear concentrations increase.

### Dorsal hubs preferentially enrich and persist at target sites

To determine whether Dorsal hubs preferentially form at transcriptionally active target sites, we performed two-color imaging of Dorsal-mNeonGreen with MCP-mCherry to visualize the site of transcription from MS2 reporters. We characterized hub formation at *snail (sna*) and *short gastrulation (sog*) target loci, as well as *hunchback (hb*) as a non-Dorsal target control (Figs. 2A-B, S2, Movies S4, S5).

**Figure 2:**
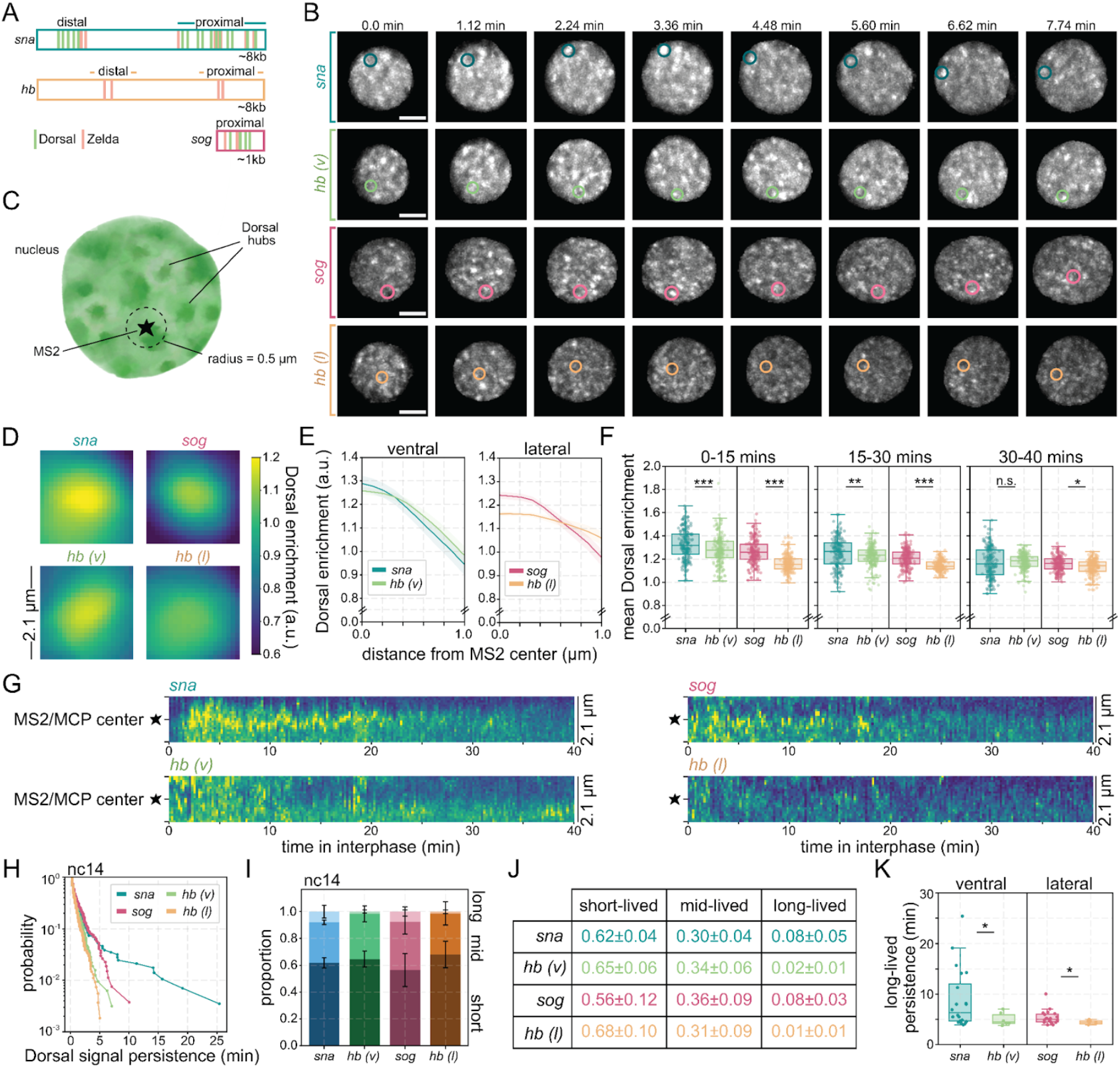
Dorsal accumulations are persistent at target genes and transient at non-targets. **(A)** Schematic of *sna, sog*, and *hb* enhancers with Dorsal and Zelda (a cofactor common to all three enhancers) binding sites shown. Each enhancer was positioned directly upstream of its promoter, followed by 24 MS2 sequences, and the *yellow* coding sequence (see Table S1 for genotypes of embryo imaged). Dorsal target genes, *snail* and *sog*, are expressed along the ventral and lateral surfaces respectively while *hunchback* is expressed anteriorly. *hunchback* is not a Dorsal target gene and is used as a negative control for both the anterior-ventral surface (*hb(v)*) to compare to *snail* or on the anterior-lateral surface (*hb(l)*) to compare to *sog*. **(B)** Representative maximum intensity projections of single nuclei with Dorsal-mNeonGreen over ∼8 mins of nc14. The MS2/MCP reporter location is circled in each image. The cytoplasmic Dorsal signal is masked, and contrast was adjusted independently for each row to aid visualization. Scale bars are 2 microns. **(C)** Schematic of analysis, a single nucleus with Dorsal hubs is depicted,the MS2/MCP spot is denoted as a star. A 0.5-micron radius sphere is considered around the MS2/MCP spot for quantification of Dorsal signal at the reporter. **(D)** Images show average Dorsal enrichment centered at the MS2/MCP site. Images are 2.1 µm x 2.1 µm. The color bar shows the level of Dorsal enrichment in the image (Dorsal intensity over nuclear mean intensity). **(E)** Average radial profile with standard error centered at MS2/MCP spots shows Dorsal enrichment as a function of distance from MS2/MCP spot center. **(F)** Boxplots showing the distribution of mean Dorsal enrichment within 0.5 micron of the MS2/MCP spot during the beginning, middle, and end of nc14 at each reporter. A Mann-Whitney U-test was performed to determine significance and following p-values were used: *p<0.05, **p<0.01, and ***p<0.001 **(G)** Representative kymographs (one nucleus per reporter is shown) shows the Dorsal enrichment centered at the MS2/MCP spot across time in nc14. **(H)** Survival plots showing the length of time any hub is consistently present within the interaction sphere for each reporter for nc14. **(I)** Stacked bar plot showing the relative proportions of short-, mid-, and long-lived Dorsal enrichment events as classified by a three-component Gaussian Mixture Model across target and non-target loci. **(J)** Table listing the mean percentages and standard deviations for each population shown in (I). **(K)** Box plots show the average duration of persistent hub presence within the long-lived category. Each persistence event from 3 embryo replicates with 5 nuclei per embryo for each reporter are plotted. A Mann-Whitney U-test was performed to determine significance and following p-values were used: *p<0.05, **p<0.01, and ***p<0.001.

Both *sna* and *hb* were analyzed using BAC reporters (*42*) with *sna* expressed in ventral nuclei and *hunchback (hb*) as a non-Dorsal target control in both ventral (*hb(v)*) and lateral (*hb(l)*) nuclei. Conversely, *sog* was driven by a transgenic reporter within its lateral expression domain (*43*). These reporters differ substantially in the number and arrangement of Dorsal binding sites with *sna* containing the highest site density (∼16 sites) compared to *sog* (∼4 sites) and *hb* (no Dorsal binding sites) (Figs. 2A, S2). Importantly, all constructs were integrated into the same genomic docking site on chromosome III, allowing us to examine enhancer grammar as the primary variable. Visually, we found that Dorsal hubs are most stably associated with *sna* and slightly less so at *sog* compared to non-specific interactions at *hb(v*) and *hb(l*) (Movies S4, S5).

We built an analysis pipeline to localize and track sites of nascent transcription from the MS2/MCP signal over time and the kinetics of Dorsal interactions at these sites (Fig. 2C, Movie S6). First, to quantify site-specific enrichment independently of hub segmentation, we measured the average Dorsal intensity centered on the MS2/MCP spot over time (Fig. 2D–E). Due to the higher nuclear background in ventral nuclei, the hub enrichment is higher at *hb(v*) than *sog*. However, when comparing *hb(v*) with *sna*, and *hb(l*) with *sog*, our analysis shows that Dorsal is enriched at actively transcribing target genes relative to non-target controls. The radial profiles demonstrate that Dorsal enrichment at target loci decays substantially within ∼0.5 µm of the MS2/MCP site with *sog* enrichment approaching *hb(l*) levels at this distance (Fig. 2E). We therefore used a 0.5 µm sphere average to capture specific locus-associated signals while reducing noise and accounting for slight spatial offsets of hubs relative to the MS2/MCP spot (Fig. 2F). The mean enrichment in this sphere closely matches the trends of enrichment measured directly at the MS2/MCP center. Consistent with the nucleus wide trends (Fig. 1), Dorsal enrichment at MS2/MCP correlates strongly linearly with nuclear Dorsal concentration (Figs. S3A), suggesting that hubs persistently form at target genes and the concentration of Dorsal within them scales with available nuclear Dorsal. Towards the end of nc14 as the Dorsal nuclear concentration drops, so does hub intensity, and the distinction between target and non-target loci is diminished.

Examination of kymographs centered on MS2/MCP spots (Fig. 2G) show that target loci exhibit extended periods of sustained enrichment compared to non-target loci. To characterize the temporal stability of hub presence at a locus, we defined a “persistent enrichment event” as a continuous time interval when the mean Dorsal intensity within 0.5 microns of the center of the MS2/MCP spot exceeds the nuclear mean intensity for at least three consecutive frames. We observed a wide distribution of persistence times at each reporter locus. Survival probability analysis of persistent enrichment times shows a significantly increased long-lived population (i.e. stable hub formation) at target loci versus non-targets (Fig. 2H). To further characterize persistence, we fit the distribution enrichment-segment durations to a three-component Gaussian mixture model (Fig. S3B) and classified segments as short-, mid-, and long-lived. Both *sna* and *sog* exhibit rare but pronounced long-lived enrichment segments (∼8% of events in nc14) (Fig. 2I–J, Fig. S3C), consistent with prior literature reporting infrequent but stable hub associations to genomic sites (*16, 22*). However, the long-lived population is significantly more persistent at *sna* than at *sog* suggesting site-specific or nuclear concentration dependent effects (Fig. 2K).

Since the experiments described above were performed using a transgenic reporter inserted in the same genomic location, to assess differences at endogenous gene loci we repeated these experiments with the same imaging conditions but now using MS2 knock-in *sna* and *sog* fly lines containing 24 MS2 stem loops placed in the 3’ or 5’ untranslated region at each endogenous gene locus (*44*) (Fig. S3D–K, Movie S7). Due to *sog* being on the X chromosome, we solely imaged female embryos while *sna* embryos were not separated by sex for analysis. In these endogenous knock-ins, relative enrichment and elevated persistence were preserved. Notably, endogenous *sna* exhibited a markedly higher proportion of long-lived enrichment segments (∼40%) (Fig. S3K). Absolute enrichment values and persistence differed slightly from transgenic reporters. These differences could be due to several factors. For instance, in the case of *sog* the endogenous locus contains additional enhancers, and for both genes, the chromatin context is different than that of the transgenic reporters inserted in a gene desert. Additionally, these genes have different nuclear positions (e.g. *sna* tends to be apically positioned (*45*)) that may affect the ability to call specific hub localization due to variation in surrounding genomic site density. For example, lower surrounding site density could lead to enhanced hub detection near the nuclear surface thus enhancing enrichment and persistence measurements. Despite these differing values, quantifying enrichment and persistence in the endogenous condition still leads to the conclusion that Dorsal hub properties are specific and tunable at different target loci. However, due to the different concentrations within nuclei that express *sog* and *sna*, absolute persistence values cannot be directly compared between these loci; therefore, we next leveraged the natural concentration gradient within the *sog* expression domain to isolate the effects of nucleoplasmic background.

To assess the effects of Dorsal nuclear concentration, we imaged the *sog* reporter near the dorsal–lateral transition point where Dorsal concentration is relatively low (as in Fig. S1A–B) and compared them to more ventrally positioned nuclei that remain within the *sog* domain but have higher nuclear concentrations. We refer to these populations as *sog*^*lateral*^ and *sog*^*medial*^ respectively (Fig. S4A–B, Movie S8). We observed clear differences in Dorsal enrichment relative to the nuclear mean between these two populations (Fig. S4C– E). In medial nuclei where nucleoplasmic Dorsal concentration is higher, the relative enrichment at the locus appears reduced. This reduction reflects an increased background intensity, since as nuclear background increases, the contrast between locus-associated enrichment and the surrounding nucleoplasm diminishes. Persistence analysis revealed a reduced long-lived enrichment population in *sog*^*medial*^ nuclei compared to *sog*^*lateral*^ nuclei (Fig. S4F–I). This reduction in persistence does not necessarily indicate reduced binding-site occupancy. In fact, previous work has shown that for another Dorsal target gene, *zen*, Dorsal maintains similar levels of occupancy in lateral and ventral regions despite changes in nuclear concentration (*46*). Therefore, the change in persistence at *sog* can be attributed to elevated background, where enrichment segments are more likely to fall below the nuclear mean threshold, shortening the apparent persistence of above-background enrichment events. Our persistence measurements thus represent a conservative estimate of enhancer-specific differences in hub stability.

Together, these data indicate that Dorsal enrichment and persistence is site-specific. While Dorsal hubs transiently appear to associate with non-target loci, these interactions are likely incidental, and stable hub enrichment is restricted to target genes.

### Hub enrichment and persistence do not strongly predict burst parameters

To assess whether Dorsal hub interactions are quantitatively linked to transcriptional kinetics, we analyzed the relationship between hub properties and transcriptional burst kinetics (Fig. 3A). Transcription occurs in bursts corresponding to transitions between inactive and active promoter states (*47*). We segmented MS2/MCP fluorescence traces normalized to MCP background levels into individual bursts by identifying changes in the rate of increase of MS2/MCP signal (see Methods) (Fig. 3B). For each reporter, we extracted burst parameters including amplitude (amount of transcripts), loading rate (slope of rise), total output (integrated intensity), duration of bursts, and frequency (interval between bursts) (*48*). We then quantified three hub-related features at each locus: (1) Dorsal intensity and (2) Dorsal enrichment at the MS2/MCP site at the burst onset, and (3) the fraction of the burst duration with persistent Dorsal enrichment (Figs. 3C, S5).

**Figure 3:**
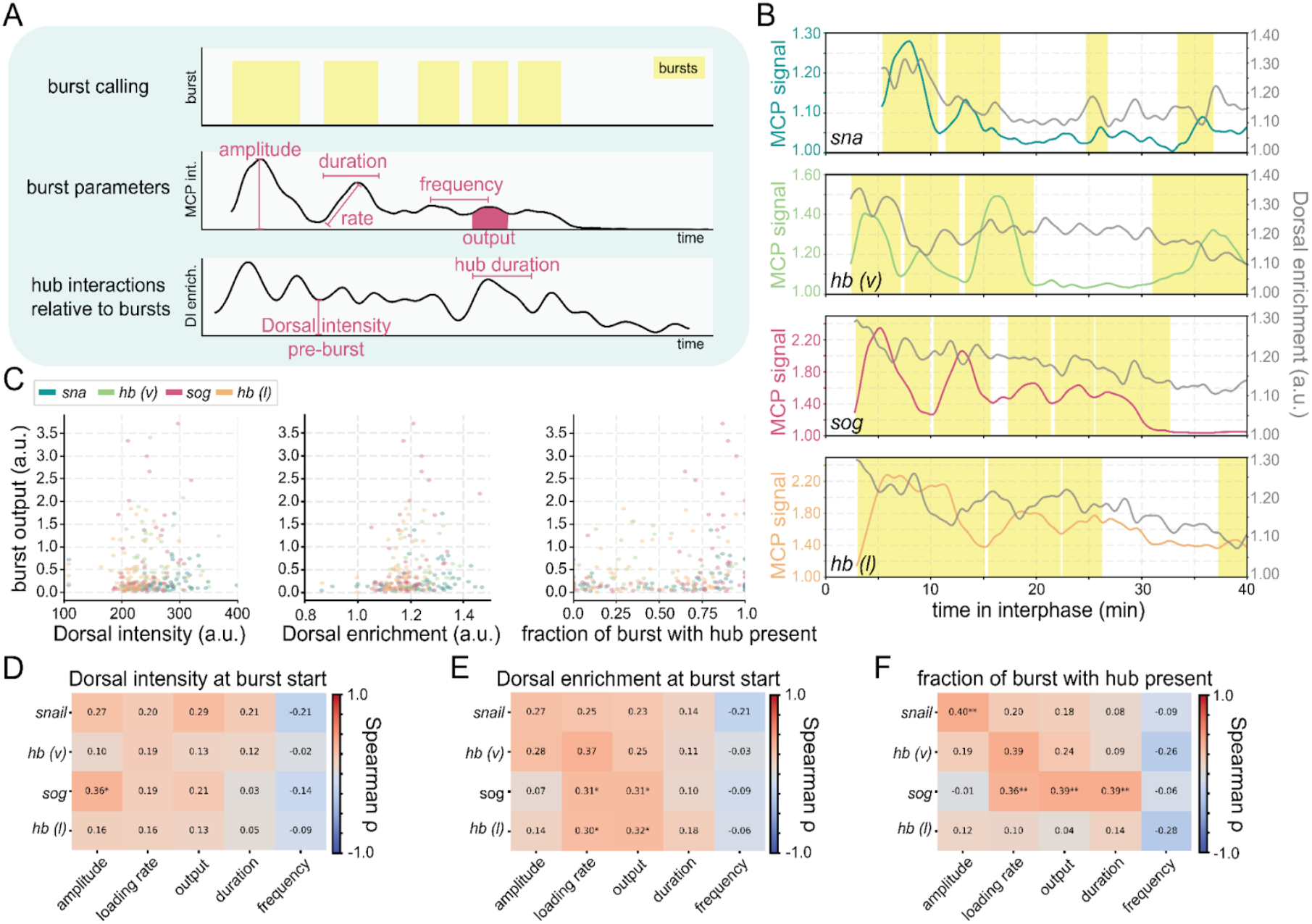
Dorsal hubs show minimal correlation with transcriptional burst kinetics. **(A)** Schematic showing the burst calling, burst parameters extracted from the MS2/MCP intensity, and the hub interaction parameters quantified from the Dorsal hub overlap and intensity. Bursts are identified in yellow. **(B)** Representative traces of MS2/MCP intensity (colored) and Dorsal intensity (grey) at each reporter gene in nc14. **(C)** Scatter plots showing burst output with the mean Dorsal intensity before the burst (left), the mean Dorsal enrichment before the burst (center), or the fraction of time during a burst when a hub is persisting near the MS2/MCP site (right). Marker colors correspond to each reporter analyzed. **(D-F)** Heat maps show the Spearman’s rank correlation coefficient (ρ) between the Dorsal intensity at burst start **(D)**, Dorsal enrichment at burst start **(E)**, and fraction of burst with hub present **(F)** for each gene and each burst parameter. The asterisks indicate p-values: *p<0.05, **p<0.01, and ***p<0.001.

Across target and non-target reporters, correlations between hub metrics and burst parameters were minimal (Fig. 3D– F). Because the MS2/MCP signal is detectable only after transcription initiation, these analyses are restricted to periods of active transcription and do not capture hub behavior preceding burst onset. The physical position of the locus cannot be reliably tracked during extended transcriptional off-intervals. While the locus position could be interpolated between active bursts, the rapid, independent three-dimensional motion of the chromatin locus risks significant spatial misalignment. Such interpolation errors could lead to false positives or negatives when calling local Dorsal enrichment or tracking hub persistence. Therefore, to ensure data integrity, metrics requiring accurate off-time tracking were excluded from this analysis. It remains a distinct possibility that hub dynamics occurring prior to or inbetween a transcriptional burst are key drivers of burst parameters or that off-times or intervals between bursts could be correlated with hub properties. Nevertheless, our results indicate that while Dorsal hubs are enriched at target loci, variation in hub intensity or persistence at a given enhancer at the start of or during a transcription burst does not robustly predict burst parameters.

### Reducing Dorsal motifs within *snail* decreases hub enrichment and persistence

To test whether enhancer binding-site number directly modulates Dorsal hub properties, we leveraged the dual-enhancer structure of the *snail* locus. We focused on this locus to remove the concentration effects between enhancers active in different concentration regions (like *sog* and *sna*). The full *snail* reporter (*sna*) contains both the proximal and distal enhancers (*42*), so we compared this construct to reporters containing only the proximal enhancer (*snaPE*) (gift from Bomyi Lim) or only the minimal distal enhancer (*snaDE*) (*39*), progressively reducing the total number of Dorsal binding sites (Fig. 4A–B, Movie S9). We included *hunchback* imaged on the ventral surface (*hb(v)*) as a negative control. Quantification of Dorsal enrichment at the MS2/MCP site revealed a reduction in enrichment from *sna* to either *snaPE* or *snaDE* (Fig. 4C–E). Notably, enrichment of both *snaPE* and *snaDE* approached levels indistinguishable from *hb(v*) suggesting that a minimum density of binding sites may be required to generate enrichment detectable above nuclear background.

**Figure 4:**
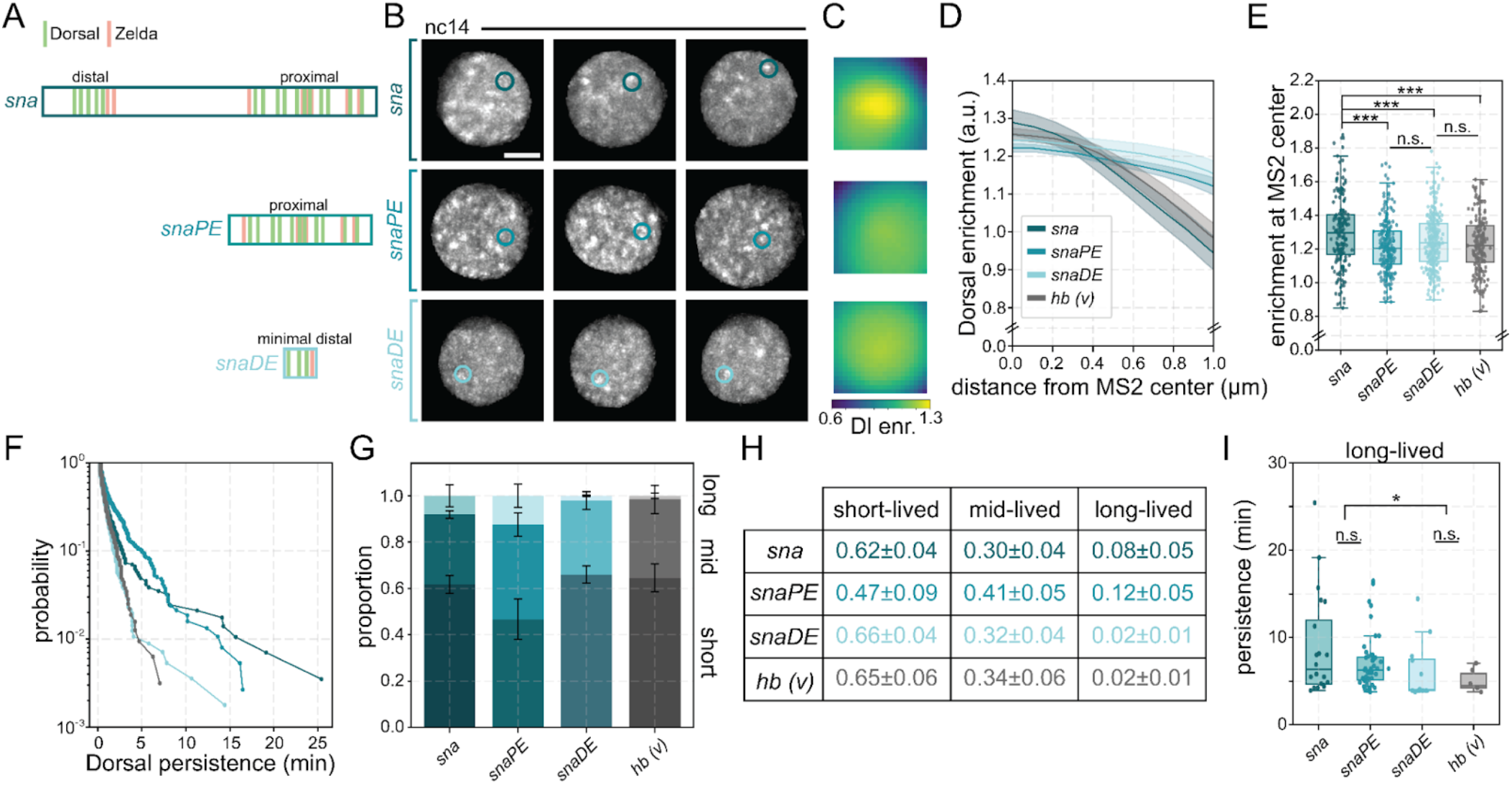
Reducing Dorsal motifs decreases hub enrichment and persistence. **(A)** Schematic depicting enhancers with Dorsal and Zelda binding sites for each *snail* reporter with either both enhancers (*sna*) (∼16 Dorsal binding sites), the proximal enhancer (*snaPE*) (∼11 Dorsal binding sites), or the distal enhancer (*snaDE*) (∼3 Dorsal binding sites). Each enhancer was positioned just upstream of its promoter, 24 MS2 sequences, and the *yellow* coding sequence (see Table S1 for genotypes of embryo imaged). Each of these *snail* reporters were imaged along the ventral surface in nc14. **(B)** Representative maximum intensity projections of single nuclei with Dorsal-mNeonGreen in nc14. The MS2/MCP spot is circled in each image. The cytoplasmic Dorsal signal is masked, and contrast was adjusted independently for each row to aid visualization. Scale bars are 2 microns. **(C)** Images show average Dorsal enrichment centered at the MS2/MCPsite. Images are 2.1 µm x 2.1 µm. The color bar on the bottom shows the level of Dorsal enrichment in the image (Dorsal intensity over nuclear mean intensity). **(D)** Average radial profile with standard error centered at the MS2/MCP spot shows Dorsal enrichment as a function of distance from MS2/MCP spot center. **(E)** Boxplots showing the distribution of the Dorsal enrichment at the center of MS2/MCP site (radius = 0.0 µm in (D)). A Mann-Whitney U-test was performed to determine significance and following p-values were used: *p<0.05, **p<0.01, and ***p<0.001. **(F)** Survival plots showing the length of time any hub is consistently present within the interaction sphere for each reporter for nc14. **(G)** The proportion of long-, mid-, and short-lived persistent hub formation in each bin is shown with error bars designating standard deviation among embryo replicates. **(H)** The proportion of short-, mid-, and long-lived hub persistence events are listed with their mean and standard deviation from (G) in the table. **(I)** Box plots showing the average duration of consistent hub interactions within the long-lived populations. Each persistence event from the following nuclei are plotted. N = *sna:* 18 nuclei in 3 embryos; *snaPE:* 22 nuclei in 3 embryos; *snaDE:* 25 nuclei in 4 embryos; and *hb(v):* 18 nuclei in 4 embryos. A Mann-Whitney U-test was performed to determine significance and following p-values were used: *p<0.05, **p<0.01, and ***p<0.001.

Although absolute levels remained near background, both *sna* and *snaPE* displayed significantly higher proportions of long-lived enrichment segments and longer persistence durations compared to *snaDE* and *hb(v*) (Fig. 4F–I). In contrast, persistence at *snaDE* was statistically indistinguishable from *hb(v*). These results indicate that increasing Dorsal binding-site number enhances both enrichment magnitude and persistence, and differences between three binding sites and the non-specific control are indistinguishable in our assay.

To further probe the sensitivity limits of our detection, we imaged additional reporters in which one or two Dorsal binding sites were removed from the distal enhancer (*49*) (*snaDE-1xdl* and *snaDE-2xdl*; Fig. S6A–B, Movie S10). Across these constructs, neither enrichment nor persistence differed significantly from the unmodified *snaDE* reporter or from *hb(v*) (Fig. S6C–H). These data indicate that further reductions in binding-site number within the distal enhancer fall below the threshold required for detectable changes in hub properties in our assay.

We next tested whether altering non-Dorsal binding sites within the enhancer influences hub behavior. Zelda is a pioneer transcription factor known to regulate early embryonic genes and to form its own nuclear hubs (*9, 39*). We compared the distal enhancer reporter to a modified reporter containing an additional Zelda binding site added 5’ (*snaDE+1zld*) (*39*) (Fig. 5A–B, Movie S11). Unexpectedly, addition of a single Zelda binding site resulted in a small but significant decrease in Dorsal enrichment at the MS2/MCP site (Fig. 5D, F, H) without altering Dorsal persistence (Fig. 5J, L; Fig. S7A–D). We then examined Zelda localization at these reporters using an endogenously tagged mNeonGreen-Zelda line (*9*) (Fig. 5C, Movie S12). Zelda enrichment increases significantly at *snaDE+1zld* relative to *snaDE* (Fig. 5E, G, I) although Zelda persistence remains unchanged (Fig. 5K, M; Fig. S7C–D). While Zelda famously acts as a pioneer factor to facilitate Dorsal recruitment globally (*50*), our data suggest that this relationship is highly sensitive to local cis-regulatory architecture. A spacing of 32 bp places the two transcription factors in a range where steric hindrance or unfavorable orientation could disrupt cooperative assembly, resulting instead in localized spatial competition. Consequently, instead of acting cooperatively, the binding of Zelda at this specific position may physically constrain local Dorsal binding, manifesting as the modest decrease in concurrent Dorsal enrichment (Fig. 5F).

**Figure 5:**
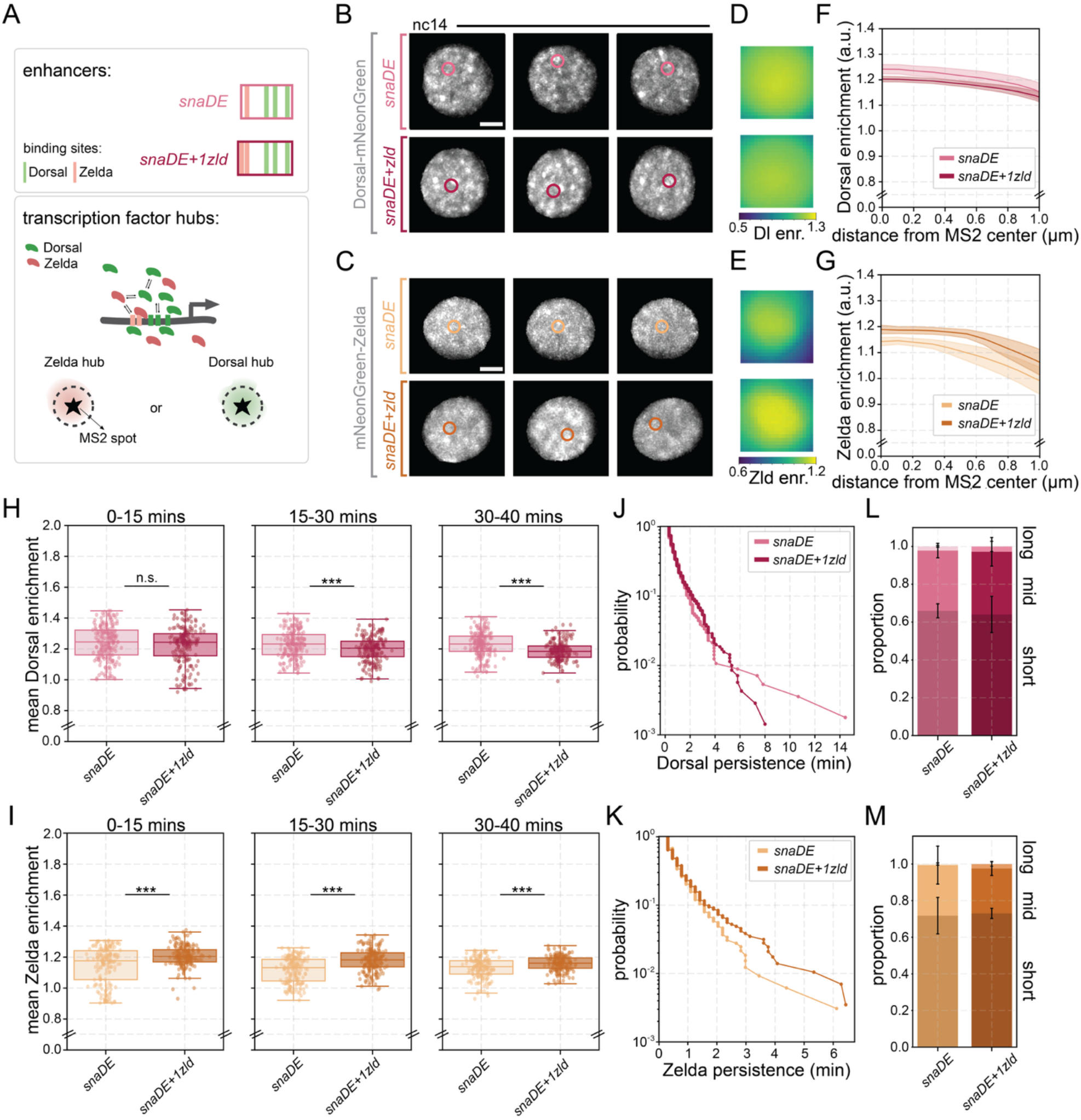
Adding a single binding site for Zelda increases Zelda enrichment at snail distal enhancer. **(A)** Schematic depicting enhancers with Dorsal and Zelda binding sites for *snail* distal enhancer reporter without (*snaDE*) or with (*snaDE+1zld*). Each enhancer was positioned just upstream of its promoter, 24 MS2 stem loops, and the *yellow* coding sequence (see Table S1 for exact genotypes of embryo imaged). Each *snail* reporter was imaged at the ventral surface in nc14. These reporters were imaged with either Dorsal-mNeonGreen or Zelda-mNeonGreen to visualize hubs formed by each TF. **(B-C)** Representative maximum intensity projections of single nuclei with Dorsal-mNeonGreen **(B)** or mNeonGreen-Zelda **(C)** in nc14. The MS2/MCP spot is circled in each image. The cytoplasmic Dorsal signal is masked, and contrast was adjusted independently for each row to aid visualization. Scale bars are 2 microns. **(D-E)** Images show average Dorsal enrichment centered at the MS2/MCP site. Images are 2.1 µm x 2.1 µm. The color bar on the bottom shows the level of Dorsal **(D)** or Zelda **(E)** enrichment in the image (local intensity over nuclear mean intensity). **(F-G)** Average radial profile with standard error centered at the MS2/MCP spot shows Dorsal **(F)** or Zelda **(G)** enrichment as a function of distance from MS2/MCP spot center. **(H, I)** Boxplots showing the distribution of mean Dorsal **(H)** or Zelda **(I)** intensities during the beginning, middle, and end of nc14 at each gene. A Mann-Whitney U-test was performed to determine significance and following p-values were used: *p<0.05, **p<0.01, and ***p<0.001. **(J, K)** Survival plots showing the length of time a Dorsal **(J)** or Zelda **(K)** hub is consistently present within 0.5 µm of MS2/MCP center for nc14. **(L, M)** Using a Gaussian Mixture Model, each Dorsal **(L)** or Zelda **(M)** hub persistence event was categorized into short-, mid-, or long-lived bins. The proportion in each bin is shown with error bars designating standard deviation among embryo replicates. N = *snaDE* (Dorsal): 25 nuclei in 4 embryos; *snaDE+1zld* (Dorsal): 36 nuclei in 4 embryos; *snaDE* (Zelda): 24 nuclei in 3 embryos; and *snaDE+1zld* (Zelda): 19 nuclei in 3 embryos.

Together, these results demonstrate that enhancer binding site composition tunes transcription factor enrichment and persistence at actively transcribing reporter sites. Dorsal hub enrichment and persistence decrease when lowering binding-site number, but these measurements are constrained by detection limits imposed by nuclear background. Moreover, addition of a single Zelda binding site is sufficient to measurably increase Zelda enrichment while modestly altering Dorsal enrichment, suggesting that enhancer composition can differentially shape factor-specific hub properties.

### Enhancer-dependent Dorsal hub properties are dictated by molecular binding kinetics

We hypothesized that if Dorsal hub properties simply reflect binding kinetics at enhancer-encoded binding sites, then a kinetic binding model should be sufficient to recapitulate our experimental results on hub enrichment and persistence. To test this, we incorporated single-molecule tracking–derived kinetic parameters into a quantitative model of transcription factor binding (*51*). To parameterize the simulations, we performed SMT experiments in live Drosophila embryos as previously shown (*8, 9, 20, 21, 41, 52*). We used a fly line containing endogenously-tagged Dorsal-mEos4a to enable sparse photoconversion for single molecule detection at both ventral and lateral surfaces of the embryo (*41*). Using long exposure times (500 msec) to blur out the unbound population of molecules and reduce the effects of photobleaching, we measured the residence time of Dorsal to be approximately four seconds in both regions (Fig. S8A–B). Using short exposure times (10 msec) to track both bound and unbound molecules, we estimated diffusion coefficients, and the immobile and free fractions (*53*) and found them to be similar (∼58%) in both ventral and lateral nuclei (Fig. S8C–D).

Using our SMT data (Fig. S8) we simulated the diffusion and binding kinetics of Dorsal molecules in nc14 nuclei. We inferred the number of Dorsal binding sites (*50*) and Dorsal concentrations (*54*) from previously published data (for a full list of parameters, see Table S2). We simulated diffraction-limited microscope images by convolving the simulated molecule positions with the microscope’s point spread function, mimicking the signal recorded by our camera sensor. These images were generated in 3D to obtain z-stacks over time, taking into account variables such as slice spacing and exposure time, to match our experimental data. We inserted binding site arrays to match the *snail* (Fig. 6) and *sog* enhancers (Fig. S9A-G) we examined experimentally. Using the same analysis pipelines we applied to our experimental data, we characterized hub enrichment and persistence in our simulated data

We found that our model is able to recapitulate the differences we see in hub properties at different *snail* enhancers (full, proximal, or distal) (Fig. 6). Hub enrichment scales with the number of Dorsal binding sites as in experimental data (Fig. 6C–E), however, the enrichment magnitude is increased relative to our real data. This increase is likely due to two factors: lower background noise in the simulated images and spread of enrichment signal in our experimental data due to our reliance on using the MS2/MCP spot (i.e. the distance of the MCP-mCherry signal plus the stem loops from the enhancer) for estimating the position of the binding sites as opposed to using the precise position of binding sites in our simulations. The persistence of Dorsal signals at different reporters also follows the same trend as our experimental data, with persistence scaling with the number of binding sites (Fig. 6H–K).

**Figure 6:**
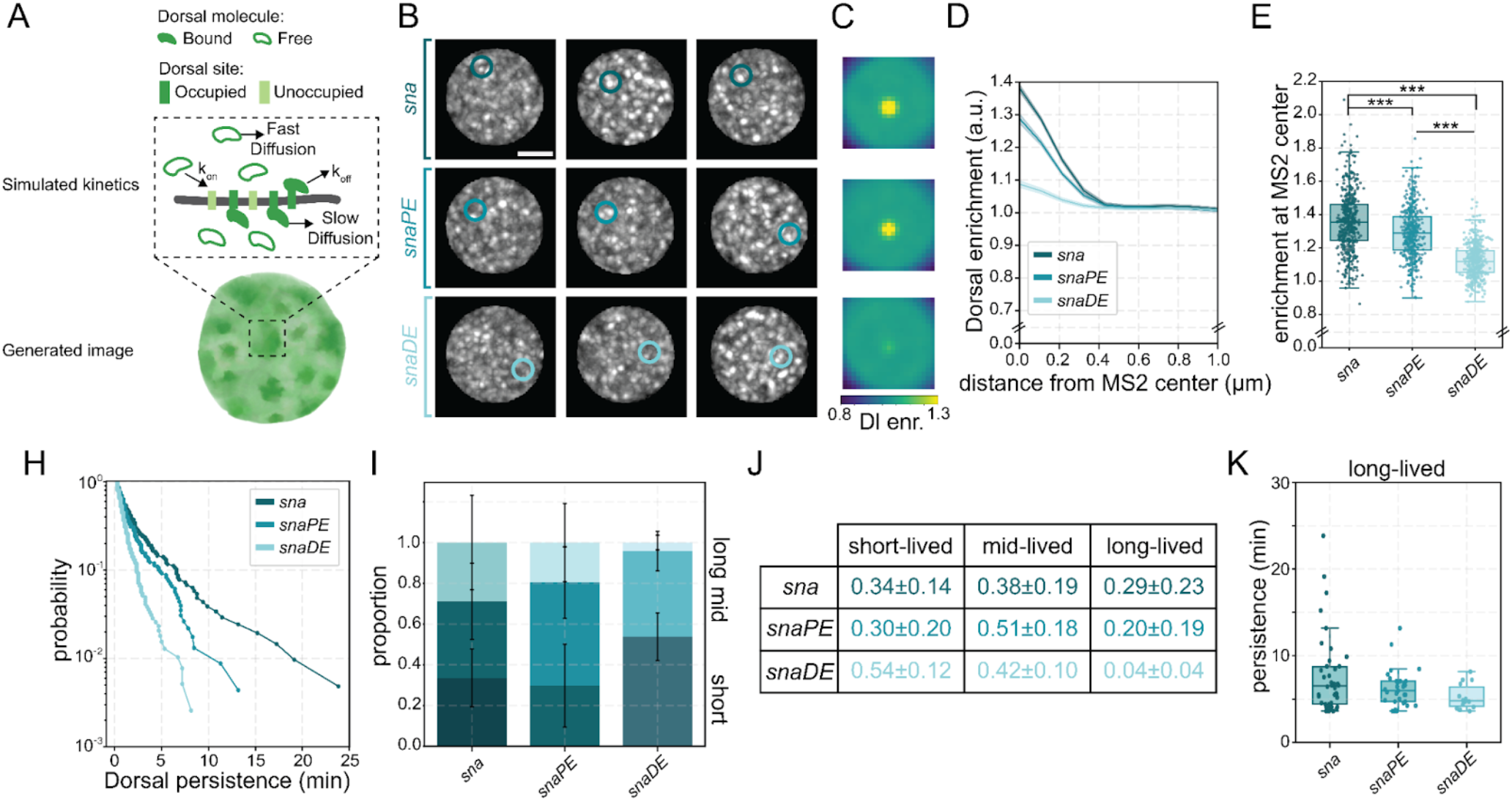
Molecular binding kinetics simulations recapitulate hub enrichment and persistence. **(A)** Schematic showing the generation of synthetic nucleus images by modeling Dorsal single-molecule kinetics and binding (*k*_*on*_ = binding probability; *k*_*off*_ = disassociation rate). **(B)** Representative maximum intensity projections of simulated single nuclei with Dorsal signal in grey. The binding site cluster is circled in each image. Contrast is scaled equally for all conditions. Scale bars are 2 microns. **(C)** Images show simulated average Dorsal enrichment centered at the binding site clusters. Images are 2.1 µm x 2.1 µm. The color bar on the bottom shows the level of Dorsal enrichment in the image (Dorsal intensity over nuclear mean intensity). **(D)** Average radial profile with standard error centered at the binding site cluster shows Dorsal enrichment as a function of distance from spot center. **(E)** Boxplots showing the distribution of the Dorsal enrichment at the center of the binding site cluster (radius = 0.0 µm). A Mann-Whitney U-test was performed to determine significance and following p-values were used: *p<0.05, **p<0.01, and ***p<0.001. **(H)** Survival plots showing the length of time any hub is consistently present within the interaction sphere for each reporter. **(I)** The proportion of long-, mid-, and short-lived persistent hub formation in each bin is shown with error bars designating standard deviation among embryo replicates. **(J)** The proportion of short-, mid-, and long-lived hub persistence events are listed with their mean and standard deviation in the table. **(K)** Box plots showing the average duration of consistent hub interactions within the long-lived populations. Each persistence event from the following nuclei are plotted. N = 15 nuclei per simulated enhancer. A Mann-Whitney U-test was performed to determine significance and following p-values were used: *p<0.05, **p<0.01, and ***p<0.001.

Notably, our simulations incorporate cooperativity by utilizing a Hill-like coefficient (*n* = 100) (*55, 56*) which dynamically scales the binding probability rather than imposing an equilibrium sigmoidal curve (see Methods). We repeated these simulations with Hill-like coefficients of 2 and 4, as reported for NF-κB (Dorsal’s mammalian homolog) (*57*) which yielded qualitatively similar results (Fig. S10 A-H). Lowering the Hill-like coefficient decreases overall enrichment and persistence, yet more clearly exposes binding-site-number scaling. Importantly, completely removing cooperativity fails to scale Dorsal persistence with site number (Fig. S10 I-V).

Additionally, we recapitulated the hub properties at *sog*^*medial*^ and *sog*^*lateral*^ and found that the increased concentration in the ‘medial’ embryonic region did lead to lower persistence consistent with our data (Fig. S9A-G). Furthermore, we used our simulation framework to illustrate the technical limits of hub detection by evaluating how background noise impacts our ability to resolve weak enrichment of *snaDE* and our negative control *hb* (Fig. S9H– N). By decreasing the simulated background noise parameter five-fold, a clear difference in enrichment and persistence emerged between *hb*^*low*^ and *snaDE*^*low*^ that was previously obscured by background fluctuations. This indicates that low-occupancy or transient hubs at weaker enhancers are not entirely absent but rather fall just below the detection threshold of our standard imaging pipeline. Therefore, our simulations demonstrate that molecular binding kinetics, and binding site number, are sufficient to explain enhancer-specific hub properties.

## Discussion

Our findings demonstrate that transcription factor hubs are not homogeneous entities but instead reflect binding kinetics at specific enhancer sequences. We show that across synthetic and endogenous reporters Dorsal hub persistence and enrichment are enhancer specific. These properties scale with the number of binding-sites and nuclear concentration but only weakly correlate with transcriptional burst kinetics (Fig. 7). These observations suggest that hubs reflect the kinetics of TF binding at enhancers rather than constituting an independent regulatory layer that determines instantaneous transcriptional output.

**Figure 7:**
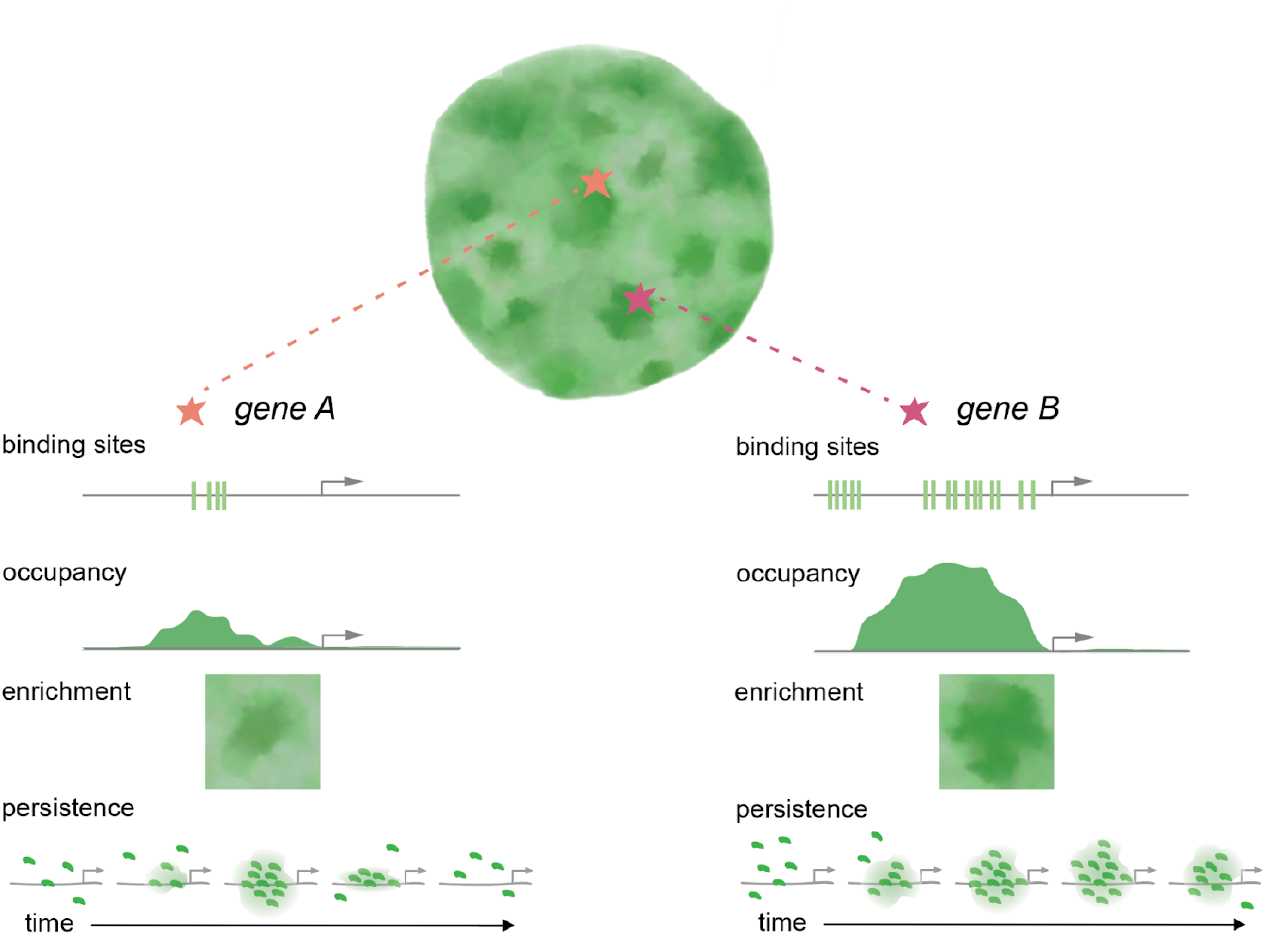
Transcription factor hub properties reflect enhancer binding site number. **(A, B)** Binding sites at *cis-*regulatory regions mediate changes in Dorsal gene occupancy which is reflected in hub properties apparent at the locus. As a result, different target sites with lower **(A)** or higher **(B)** number of binding sites have differing enrichment and persistence of Dorsal hubs.

We found that the nuclear density of Dorsal hubs does not depend on nuclear concentration, while hub enrichment does. The decoupling of density from concentration is inconsistent with models where hubs arise from concentration-threshold dependent assembly of condensates. Although Dorsal clustering has been suggested to be concentration dependent and arising from thermodynamic phase separation (*40*), our results indicate that hub density is not dictated by bulk nuclear levels. Instead, nuclear concentration appears to tune factor enrichment at existing genomic binding sites rather than creating new assemblies. This is further supported by our computational modeling which recapitulates hub properties such as enrichment and persistence through binding kinetics alone.

The cooperativity dependence required to reproduce hub persistence indicates that independent Dorsal binding is insufficient, and that local cooperative effects are important. Dorsal has been modeled with high Hill coefficients (∼100) to account for switch-like gene expression patterns (*55, 56*). This parameter could reflect local effects, where initial binding increases the probability that subsequent molecules rebind locally before escaping, or known cooperativity with cofactors such as Zelda, Snail, and Twist (*58–61*). Notably, testing moderate exponents yields qualitatively similar clustering behavior (Fig. S10A–H). Lowering the exponent decreases absolute enrichment and persistence, yet more clearly exposes binding-site-number scaling by removing the compression caused by extreme switch-like filtering. The fact that minimal cooperativity is sufficient to produce persistent hubs demonstrates that extreme, switch-like behavior is not strictly required. Crucially, whether driven by intrinsic cooperativity, local kinetic shifts, or cofactor interactions, these mechanisms remain rooted in sequence-encoded binding kinetics. Thus, phase separation-like mechanisms are not necessary to explain these observations.

By using reporters integrated in a common genomic docking site, we show that increasing Dorsal binding site density increases enrichment and persistence. Conversely, reducing binding sites progressively diminishes detectable enrichment and long-lived interactions, eventually reaching levels indistinguishable from non-specific control sites. These effects extend beyond Dorsal itself. The addition of a single Zelda binding site measurably increases Zelda enrichment while modestly reducing Dorsal enrichment, highlighting that enhancer composition differentially tunes factor-specific hub properties. Cis-regulatory element control of transcription factor hubs is not completely unprecedented. Prior work demonstrated that removing Zelda binding sites at *sog* reduced both Dorsal clustering and transcription, particularly in lateral regions where Dorsal concentration is lower (*32*). Other work also demonstrated that high-affinity Bicoid binding sites did not result in increased Bicoid clustering intensity, suggesting that clustering is not simply dictated by affinity (*38*). However, this prior study did not account for differences in number of binding sites or hub persistence, which may be a crucial distinguishing factor, akin to the differences observed between *snail* and *sog* in our data. Overall, our data suggests that enhancer grammar may play a key role in determining hub properties and function. A key implication of our results is that persistence and enrichment can be interpreted as realtime occupancy readouts. Genomic approaches such as ChIP-seq have shown that motif number and affinity correlate with transcription factor occupancy. Here, our time-averaged enrichment measurements correlate with binding-site number across genes *in vivo*, suggesting that hub enrichment reflects binding kinetics at enhancer sites.

Importantly, our data reveal that hub properties are only weakly predictive of transcription burst kinetics. In line with our data, previous studies on Gal4 clusters found no difference in burst size when the clusters were positioned closer to their target site and *in vivo* single molecule tracking of Gal4 at its target site suggests that burst duration and size are primarily determined by transcription factor dwell time and binding affinity (*35*). Although stable enrichment is more prevalent at target genes, variation in enrichment or persistence during active transcription does not strongly scale with burst kinetics. However, because our analysis is restricted to active phases of the transcription cycle, we cannot rule out the possibility that hub properties influence parameters during inactive periods such as the duration between bursts. Furthermore, our current framework does not evaluate the first passage time of RNA Polymerase II following mitosis, leaving open the possibility that hub properties prior to the initial transcription burst of nc14 are linked to downstream regulatory effects. Our data nonetheless suggest that while binding-site composition tunes occupancy, transcriptional bursting is likely influenced by steps downstream of TF recruitment.

As has been shown previously, the presence of a TF hub is not sufficient for transcription initiation (*33*). Indeed, we previously examined the effect of enhancer placement relative to the promoter (*62*). Downstream placement of the enhancer significantly dampened transcriptional output, yet we detected no change in local Dorsal enrichment between upstream and downstream configurations. This dissociation between enrichment and transcription further supports the conclusion that hub formation and enrichment are dictated by binding-site composition rather than act as transcriptional modulators at a given enhancer. While our initial observations here suggest that hub properties at burst onset are not strong predictors of burst parameters at a single locus, whether these features are directly linked remains an open question. Indeed, our work does not rule out hub contributions prior to activation or between bursts as previously reported (*22, 33, 63*). Additionally, recent work emphasizes that cooperative TF binding and exchange can stabilize high-occupancy states that underlie transcriptional bursting (*34, 64*). We similarly find in our modeling that cooperativity is required to reproduce the measured hub properties. Still, the differences in burst kinetics and transcriptional output between genes are likely dependent on the level of TF occupancy which is reflected in the larger differences in hub properties between reporters.

Our findings also highlight a critical practical consideration: hub detection is inherently constrained by nuclear background, and the contrast of the imaging modality being used. As nuclear Dorsal concentration increases, the background intensity is elevated, reducing contrast between locus-associated enrichment and the surrounding nucleoplasm without necessarily altering underlying binding kinetics at a locus. This limitation in detection could easily lead to misleading conclusions that condensates or hubs are discrete regulatory structures that highly concentrate molecules. Indeed, our recent work on RNA polymerase II (Pol II) clustering similarly demonstrates that Pol II clusters at active genes can be explained by transcription-dependent engagement and recruitment kinetics and are thus only visible at genes that sufficiently accumulate Pol II (*20*). Our current study is also limited by its focus on a single TF. Because these hubs are emergent properties driven by enhancer composition, understanding how multiple molecules work together at the locus is critical to seeing how TFs and transcriptional machinery jointly impact local occupancy. Developing multi-factor models of binding kinetics to explore how TFs interact with both DNA and one another remains an important future endeavor.

Together our results suggest the clustering of Dorsal, and other TFs, observed *in vivo* may be interpretable through the lens of binding kinetics and occupancy rather than phase separation. Second, they underscore the importance of cooperative binding and enhancer features in shaping apparent hub behavior. Third, they highlight how detection thresholds and nuclear background can influence conclusions about condensate formation. Finally, by demonstrating that enhancer perturbations produce quantifiable changes in enrichment and persistence, our study establishes that hub properties are encoded within DNA sequence. We suggest that the many reports of transcription factors and regulatory proteins forming condensates or hubs through higher-order mechanisms should be examined through the lens of molecular binding kinetics. Instead, rapid search processes may facilitate a high on-rate and frequent exchange of TF molecules at their targets, leading to the appearance of persistent hubs at gene loci despite short residence times. Future work combining enhancer dissection, kinetic modeling, and hub molecular composition analysis will further clarify how generalizable this binding and search process centric framework is across transcription factors and developmental systems.

## Acknowledgements

We thank Bomyi Lim and Mounia Lagha for fly lines and helpful discussions about this work. We also thank all members of the Mir Lab for feedback throughout this project. This paper was typeset with the bioRxiv word template by @Chrelli: www.github.com/chrelli/bioRxiv-word-template

## Funding

National Institutes of Health: DP2HD108775 (MM), R01GM139913 (HGG) and R01GM152815 (HGG)

Margaret Q Landenberger Foundation (MM)

Howard Hughes Medical Institute, Freeman Hrabowski Scholars Program (MM)

National Science Foundation Graduate Research Fellowship grant DGE-2236662 (SF) and DGE-1752814 (MAT)

Koret-UC Berkeley-Tel Aviv University Initiative in Computational Biology and Bioinformatics (HGG)

Winkler Scholar Faculty Award (HGG)

Chan Zuckerberg Initiative Grant CZIF2024-010479 (HGG)

## Author contributions

Conceptualization: MM, SF Methodology: SF, MM, AM, MK Software: SF, MK

Validation: SF, MM, MK

Formal Analysis: SF, MM, MK, LE Investigation: SF, MM, MK, LE Resources: MAT, HGG, AM

Data Curation: SF, MM, LE

Writing – Original Draft: SF, MM, MK, LE

Writing – Review & Editing: SF, MM, AM, MAT, HGG, MK, LE Visualization: SF, MK, LE

Supervision: MM

Project Administration: MM

Funding Acquisition: MM, SF, HGG, MAT

## Data, code, and materials availability

All data needed to evaluate the conclusions in the paper are present in the paper and/or the Supplementary Materials. All acquired microscopy images are available on request. Custom analysis software and select data used in this study is available on Zenodo (https://doi.org/10.5281/zenodo.18941868) and on Gitlab (https://gitlab.com/mir-lab/publications/dorsal-ms-code-compilation-2025.git).

## Materials and Methods

### Embryo Preparation for Live Imaging

For each MS2 experiment, *Drosophila* embryos were generated by crossing virgin female flies heterozygous for MCP-mCherry (III) and Dorsal-mNeon-Green to males containing the MS2 reporter of interest or males containing the endogenous MS2 knock-in. The resulting embryos were collected for live imaging. Only embryos positive for all three (Dorsal-mNeonGreen, MCP-mCherry, and MS2-gene-of-interest) were imaged. In the instance of the endogenous *sog-MS2* knock-in only, since *sog* is on the X chromosome, the active MS2/MCP spots would only be visible in female embryos as the MS2 array is only passed down via the paternal X chromosome. Thus all embryos were screened for active MS2/MCP spots in nc13. These selected females were then chosen for imaging nc14.

Details of all lines used for imaging and where they appear in this work are listed in Table S1. For single molecule tracking experiments, virgin female Dorsal-mNeonGreen; MCP-mCherry flies were crossed to male Dorsal-mEos4a flies. Flies positive for all three markers were used for imaging. Dorsal-mNeonGreen intensity was further used for assessing the position of the embryos along the dorsoventral axis; MCP-mCherry was used for assessing nuclear boundaries; and Dorsal-mEos4a was sparsely activated for single molecule detection. Embryos were collected over a 90-minute window and manually dechorionated by gently rolling them on double-sided tape using a blunt needle until the chorion was removed. Dechorionated embryos were then arranged on an agar pad before being transferred onto an adhesive heptane-acrylic spot, prepared by dissolving double-sided tape in heptane on a 25 mm diameter glass coverslip. Embryos were manually oriented on the coverslip either ventral side up (facing the objective) or gently rotated to position the dorsal–lateral transition region within the imaging plane (Fig. S1A,B).

### Annotation of Transcription Factor Binding Sites

Transcription factor (TF) binding sites within the enhancers were identified using the FIMO (Find Individual Motif Occurrences) tool (*65*) with position weight matrices (PWMs) obtained from (*66*) and the JASPAR database (*67*). The threshold for a significant motif match was set to a p-value<10^™3^. For *snaDE*, local annotation of Dorsal and Zelda motifs was cross-referenced with established cis-regulatory mappings (*49, 68*). While the *snaDE* sequence lacks the canonical CAGGTAG Zelda motif, it contains an AGGTAG motif near the 5′ end on the complementary strand capable of functional Zelda binding (*69*). This endogenous motif is preserved in the engineered *snaDE+1zld* construct, placing active Zelda sites in close proximity to each other and 32 bp away from a Dorsal binding site.

### Microscope Calibration

All imaging was performed on a Multimodal Optical Scope with Adaptive Imaging Correction (MOSAIC) (*70*). For volumetric experiments, the lattice light-sheet mode was used with the 488 nm and 589 nm laser lines. These laser beams were first expanded to a combined 2 mm diameter, passed through a half-wave plate to adjust polarization, and relayed into an acousto-optic tunable filter (AOTF) (Quanta-Tech, AA Opto Electronic) to selectively control wavelength and power modulation. The collimated beams were then expanded in one dimension using a Powell lens (Laserline Optics, Canada), and their width was adjusted and collimated using a pair of cylindrical lenses. The beam was subsequently relayed onto a grayscale spatial light modulator (Meadowlark Optics, AVR Optics) after passing through a second half-wave plate, where it was diffracted and projected onto a custom annular mask to define the minimum and maximum numerical aperture (NA) of the light-sheet while blocking undiffracted light. The light from the annulus was demagnified and relayed onto a resonant galvanometer scanner (Cambridge Technology, Novanta Photonics), conjugate to the sample plane, to mitigate shadowing artifacts and improve uniformity by introducing a slight wobble in the excitation angle. Finally, the light was projected onto a pair of galvanometer scanning mirrors (Cambridge Technology, Novanta Photonics) conjugate to the pupil plane, allowing scanning of the light-sheet along the x and z optical axes before being focused onto the sample using a Thorlabs TL20X-MPL excitation objective. Fluorescence emission was collected orthogonally by a Zeiss 20×, 1.0 NA detection objective and relayed onto a deformable mirror (ALPAO) positioned in the detection path to correct optical aberrations, as previously described (*71*). The emitted signal was then split using a Semrock Di03-R561-t3-25×36 dichroic beam splitter and directed onto two sCMOS detectors (Hamamatsu ORCA Fusion). The green emission channel (for 488 nm excitation) was filtered using a Semrock FF03-525/50-25 emission filter and a Chroma ZET488NF notch filter to reject laser light, while the red emission channel (for 589 nm excitation) was filtered using a Semrock FF01-615/24 emission filter and a Chroma ZET594NF notch filter

### Imaging Conditions

All volumetric imaging imaging used a multi-bessel lattice sheet with maximum numerical aperture to minimum numerical aperture ratio of 0.4/0.3 488 nm (used to excite mNeonGreen) and 589 nm (used to excite mCherry). Two color channels were acquired sequentially for a volume of 18.9 µm sampling z-planes spaced 0.3 µm apart with an exposure time of 30 msec. A laser power of 0.124 mW was used for the 488 nm laser and 1.344 mW for the 589 nm laser. Each volume was acquired four seconds after the acquisition ended of the previous stack for a total time of ∼9.56 seconds in between stacks. These conditions were kept constant for each imaging experiment aside from the single molecule tracking experiments.

For single-molecule imaging, a Gaussian light sheet was used on the MOSAIC. Briefly, after passing through the AOTF, the beam was expanded using a Powell lens then relayed onto a custom mask for filtering. The filtered sheet was then projected onto a resonant galvanometer (Cambridge Technology, Novanta Photonics). Lastly, an excitation objective (Thorlabs, TL20X-MPL) was used to focus the light-sheet onto the sample. The detection path for the emitted light is the same as described above for the lattice light-sheet configuration. For this light path, the SLM was bypassed to maximize laser power at the excitation objective. For single molecule imaging, the red emission channel (for 561 excitation) was filtered using a Semrock FF01-585/29 emission filter and a Chroma ZET561NF Notch filter. The 405 nm laser line was kept on constantly during the acquisition period for photoswitching. Data was acquired at 10 msec and 500 msec exposure times for the fast and slow single-molecule imaging experiments respectively. The excitation laser power was optimized empirically for each exposure time to achieve sufficient contrast for single-molecule tracking. The powers of the photoswitching laser were also optimized empirically to achieve low enough detection densities to enable tracking. For fast single-molecule imaging experiments, 8000 frames were acquired with 24.1 mW of the 560 laser, 123 µW of the 488 laser, and 7.3 µW of the photoswitching laser for a total acquisition time of 80 sec. For slow single-molecule imaging experiments, 400 frames were acquired with 0.66 mW of the 560 laser. 123 µW of the 488 laser. and 0.46 µW of the photoswitching laser for a total acquisition time of 200 sec. Powers were measured at the back focal plane of the excitation objective.

### Volumetric Image Analysis

All volumetric imaging datasets were pre-processed before downstream analysis using GPU-accelerated 3D deconvolution via CUDA (*72*). Each dataset was deconvolved using a Richardson-Lucy based algorithm with 8 iterations, utilizing point spread functions (PSFs) captured from bead images acquired on the day of imaging to ensure accurate correction. For visualization and rendering, we employed a custom Imaris converter, leveraging fast TIFF and Zarr file readers to efficiently generate Imaris files (*73*). Following deconvolution, images underwent nuclear segmentation using a custom-trained model in Cellpose 2.0 (*74*). Ground truth training data was generated through a combination of micro-SAM (*75*) and manual curation on MCP-mCherry images. The trained Cellpose model was then applied to segment individual slices across each dataset. To improve segmentation accuracy, a post-processing pipeline was developed to stitch segmented slices together and interpolate missing nuclear structures. We then implemented a nearest-neighbor tracking algorithm to follow nuclei across interphase in each nuclear cycle.

To quantify hub properties, we used a custom image analysis pipeline (*20, 21*) . First, nuclear intensity was normalized to its mean value to assess local transcription factor enrichments above background levels. Hubs were segmented by first applying a median filter to reduce noise, followed by image erosion and reconstruction to enhance feature contrast. The reconstructed image was then subtracted from the median-filtered image to generate a binary mask identifying high-density regions. To resolve closely associated hubs, local maxima peaks were identified and used as markers for water-shed-based segmentation, ensuring separation of fused or amorphous structures. Finally, region properties were extracted to quantify hub features, including integrated intensity, mean intensity, and size.

### MS2 Tracking

After segmenting nuclei and hubs, we manually select nuclei for further analysis under the criteria of checking if anomalies are present in the segmentation and to check if the MS2 spot is closer than 0.2 µm to the edge of the nucleus which may skew analysis. We then generate 4D images of cropped single nuclei. These selected nuclei are then subjected to another segmentation regime this time to detect the MS2/MCP spot. This regime uses a median filter, difference-of-gaussians filter, and then a percentile threshold to segment high intensity spots. The exact sigma values and percentile used were adjusted nucleus-to-nucleus to ensure the best possible MS2/MCP segmentation. Then we identify the spot inside the nucleus with the highest intensities to be the likely MS2/MCP spot. We then used a nearest neighbour algorithm to track the MS2/MCP spot in time while also manually correcting any tracking errors in each nucleus. Due to the bursty nature of transcription, we interpolate the MS2/MCP position in between bursts of visible MS2/MCP spots and extrapolate the MS2/MCP position 10 frames before the first appearance of the MS2/MCP signal while correcting for the motion and size of the nucleus.

We calculated enrichment (Dorsal intensity at a locus over the nuclear background) as previously described in (*76*). However, instead of solely using spot detection, we used the calculated MS2/MCP centroid positions tracked over time as the center position for our cropped images. We then quantified enrichment segments for our persistence measurements at the MS2/MCP spot by assessing the overlap between the hub labels and a 0.5 µm radius sphere centered at the MS2/MCP weighted centroid coordinates as we have done previously (*21*). We could then monitor the duration, intensity, and volume overlap at each locus regardless of transcriptional activity. Hub persistence durations were fit using a three-component Gaussian mixture model in order to establish short-, mid-, and long-lived populations. To rigorously determine the optimal number of populations, we evaluated model performance across a range of components: a three-component model was selected as it minimized both the Akaike Information Criterion and Bayesian Information Criterion.

### Burst Analysis

To calculate the intensity at the MS2/MCP spot, we averaged the MCP intensity within 0.5 µm radius of the MS2/MCP centroid position. Since we used MCP tagged with the mCherry fluorophore, we checked for significant signs of photobleaching. Under our imaging conditions, nuclear MCP-mCherry retains approximately 77% of its initial intensity by the end of the 40-minute imaging window (Fig. S5A). Crucially, we find that both the MS2/MCP spot and the nuclear background levels of MCP-mCherry bleach at equivalent, proportional rates. This uniform decay preserves a stable signal-to-noise ratio throughout the nuclear cycle. To minimize the technical impact of photobleaching and avoid relying on absolute intensity values, we normalize the MS2/MCP spot intensity directly to the nuclear background. Because the signal-to-noise ratio is maintained, this level of signal decay is minimal, and we consider this to be minimal bleaching allowing us to continue with our analysis.

To analyze transcriptional bursting, MS2/MCP trajectories during nc 14 were standardized to a 0–40 min timeline and smoothed using a one-dimensional Gaussian filter (σ=2 frames) with background MCP signal prior to activation masked. The initial activation burst was defined by locating the peak within a 10-minute window post-onset, with the burst end mapped to the first local minimum after the signal fell below a 50% relative half-height threshold. Subsequent bursts were identified using scipy.signal.find_peaks (minimum prominence of 0.03, minimum height of 0.05) and bounded by a localized ramp-strength check requiring a consecutive positive slope of ≥0.003 MCP units/frame over ≥7 frames. For each segmented burst spanning, we extracted three parameters: (1) amplitude, defined as the peak value of the smoothed MCP normalized intensity; (2) duration, calculated as the elapsed time of burst; and (3) total output, calculated via trapezoidal integration of the baseline-subtracted MCP curve over the burst duration.

Based on the start times of these bursts we could then assess if a hub was present at the start of a burst, the arrival time of that hub overlapping with the MS2/MCP region and then overlap intensity and volume. To assess if a hub was present, we used two different metrics: we first assessed if any hub voxels overlapped with the MS2/MCP region and then assessed if a hub was present and was over a threshold intensity value (the nuclear mean intensity).

### Fast Single Molecule Tracking (SMT) Analysis

To process the fast single-molecule tracking (SMT) data, we first removed the initial ∼1,000 frames affected by fluorophore bleaching. Maximum projections were generated and input into the Cellpose-SAM model (*77*) to obtain nuclear masks. These masks were manually edited to ensure localizations along the nuclear border were excluded. Localization and tracking of the single molecule trajectories was performed using the open source software package *quot* (*78*). For the 10 msec data the following key parameters on quot were used: (i) for detection - method = “identity”, Spot detection settings = “llr”, *k* = 1.3, *w* = 15, *t* = 18, (ii) for localization - method = “ls_int_gaussian”, window size = 9, sigma = 1.0, ridge = 0.0001, maximum iterations = 10, damp = 0.3, camera gain = 1, camera background = 108 (iii) for tracking - method = “euclidean”, pixel size (in µm) = 0.108, search radius = 0.7 µm, maximum blinking = 0 frames. We then filtered the list of tracked molecules output by quot using the nuclear masks as described earlier to ensure we analyzed only trajectories localized in the nucleus.

To analyze the fast single molecule trajectories to infer the diffusion kinetics we used Spot-On (*53*). Spot-On is a publicly available package that fits a kinetic model to the jump length distribution of the trajectories and estimates the fraction bound and diffusion coefficients for either a two state or a three state model, correcting for the probability of molecules diffusing out of focus. Trajectories from all videos were concatenated into one file per gradient position. For the analysis, we used the following parameters altered from default:

TimeGap = 12

dZ = 0.700

GapsAllowed = 0

TimePoints = 4

UseEntireTraj = 1

JumpsToConsider = 0

NumberOfStates = 2

FitIterations = 2

LocError = 0.030

UseWeights = 1

D_Free_2State = [0.2656 25]

D_Bound_2State = [0.0001 0.2656]

Single molecule trajectories were taken from a minimum of three independent embryos for all positions and imaging conditions. Detailed trajectory number and the number of videos from which the trajectories were compiled can be found in each figure caption.

### Slow Single Molecule Tracking (SMT) Analysis for Residence Times

To process the slow SMT data, the same methodology was followed as described above for fast SMT, with the exception of two *quot* parameters: *k* =1.5 and *t*=15.5. Residence times were calculated from single-molecule data acquired at 500 msec exposure times as described previously (*9, 79*). In brief, imaging at 500 msec exposure times effectively blurs out fast moving molecules, leaving the slowly diffusing and bound molecules for tracking. The longer exposure time enables lower excitation laser power, leading to less photobleaching and longer trajectories. The persistence of trajectories over time is used to calculate a survival probability, which is 1-cumulative distribution function of trajectory lengths (in number of frames), providing an estimate of a genome average residence time. An objective threshold of two seconds (*9*) was applied to the minimum trajectory length to remove the effects of tracking errors and slowly diffusing molecules. Probabilities below 10^-3^ were not considered for fitting to avoid using the sparsely populated distribution tails.

The survival probability curves were then fit to a double exponential model of the form *F *exp(-k*_*ns*_ *\*t) + (1 - F) * exp(-k*_*s*_ *\* t*) using the curve_fit function in Python where *k*_*s*_ is the slower off-rate and *k*_*ns*_ is the faster/nonspecific off-rate. An exponential weighting function was used to ensure proper estimation of the slower off-rate so that the fit was not dominated by the more numerous early-time point trajectories. As the inferred off-rate *k*_*s*_ is biased by photobleaching and chromatin motion, bias correction was performed as previously described: *k*_*s,true*_*=k*_*s*_*-k*_*bias*_, where *k*_*bias*_ is the slower-off rate estimate from fitting the survival probabilities from H2B-mEos4a trajectories imaged in the same conditions. The genome wide residence times were then calculated as *1/k*_*s,true*_. Errors reported were estimated from the standard deviations of the fit parameters using standard propagation of error methods.

### Simulations

Simulations modeling Dorsal diffusion and binding were developed in Python. Nuclei in nc14 were modeled as spheres of 5 µm and Dorsal molecules were initialized at random positions within the spheres. The concentration of Dorsal along the gradient was estimated using the known ventral concentration, 500 nM (*54*), and nuclear mean intensity ratios from our live imaging data. Dorsal binding sites are also distributed in clusters of varying densities. Each binding site is assigned a residence time with probabilities broadly based on the survival curves in Fig. S8A, with a greater proportion of sites having shorter residence times. A three-state model was used with Dorsal molecules being either freely-diffusing, specifically bound or non-specifically bound. Free diffusion was modeled by Brownian motion with a simulation time step of 10 ms to mimic our single-molecule tracking. At each time step, molecules within a 50 nm capture radius of an unoccupied inding site can bind the site with a certain probability *k*_*on*_, reflecting the accessibility of the site. Once bound, the molecule diffuses with the binding site, which undergoes fractional Brownian motion to mimic the sub-diffusive motion of DNA (Rouse polymer model). This is done via the Davies-Harte algorithm, on top of which a Gaussian jiggle is added to represent local fluctuations. For memory efficiency, fractional Brownian motion increments are generated per display frame, and then bridged together such that subsequent frames are conditioned on previous ones. After the residence time of the binding site, the molecule unbinds and returns to the freely-diffusing state.

To account for local cooperative interactions without enforcing an equilibrium sigmoidal response, we modeled cooperativity as a dynamic, occupancy-dependent increase in binding probability (*k*_*on*_). When a site in a cluster is evaluated for binding, the effective binding probability *k’*_*on*_ is scaled according to current local occupancy: *k’*_*on*_ *= k*_*on*_ *\* (1 + (S*_*coop*_ *\* N*_*occupied*_)^*n*^) where *k*_*on*_ is the baseline binding probability, *S*_*coop*_ is the cooperativity strength parameter (set to 1.0), *N*_*occupied*_ is the number of currently occupied binding sites within the cluster of binding sites, and *n* is a cooperative scaling exponent. Rather than a classical equilibrium Hill coefficient, which dictates fractional occupancy output, the exponent *n* acts as a kinetic modulation parameter that increases the *k*_*on*_ as bound molecules accumulate. In short, when cooperativity is enabled, the presence of a bound molecule within a cluster of binding sites increases the *k*_*on*_ for subsequent molecules based on n which we term a “Hill-like” coefficient.

Simulated z-stacks were recorded at intervals of 10 s with a camera exposure time of 30 ms. The molecules intensities at each simulated step within the exposure time are added together and projected on a 3D voxel grid, with an xy-spacing of 108 nm (corresponding to the pixel size on the MOSAIC camera) and a z-spacing of 300 nm (corresponding to the width of our z-stacks). Each molecule is convolved with a PSF, an anisotropic Gaussian with lateral full-width at half maximum (FWHM) of 250 nm and axial FWHM of 600 nm, the estimated post-deconvolution blur obtained by deconvolving a fiducial image from the MOSAIC with a theoretical PSF. We additionally incorporated the background noise from the camera (100 ± 5 a.u.). More about these simulations and an user-friendly interactive tool to experiment on other transcription factors can be found here (*51*).

