## Supplemental Materials for "Enhancer binding kinetics explain transcription factor hub formation"

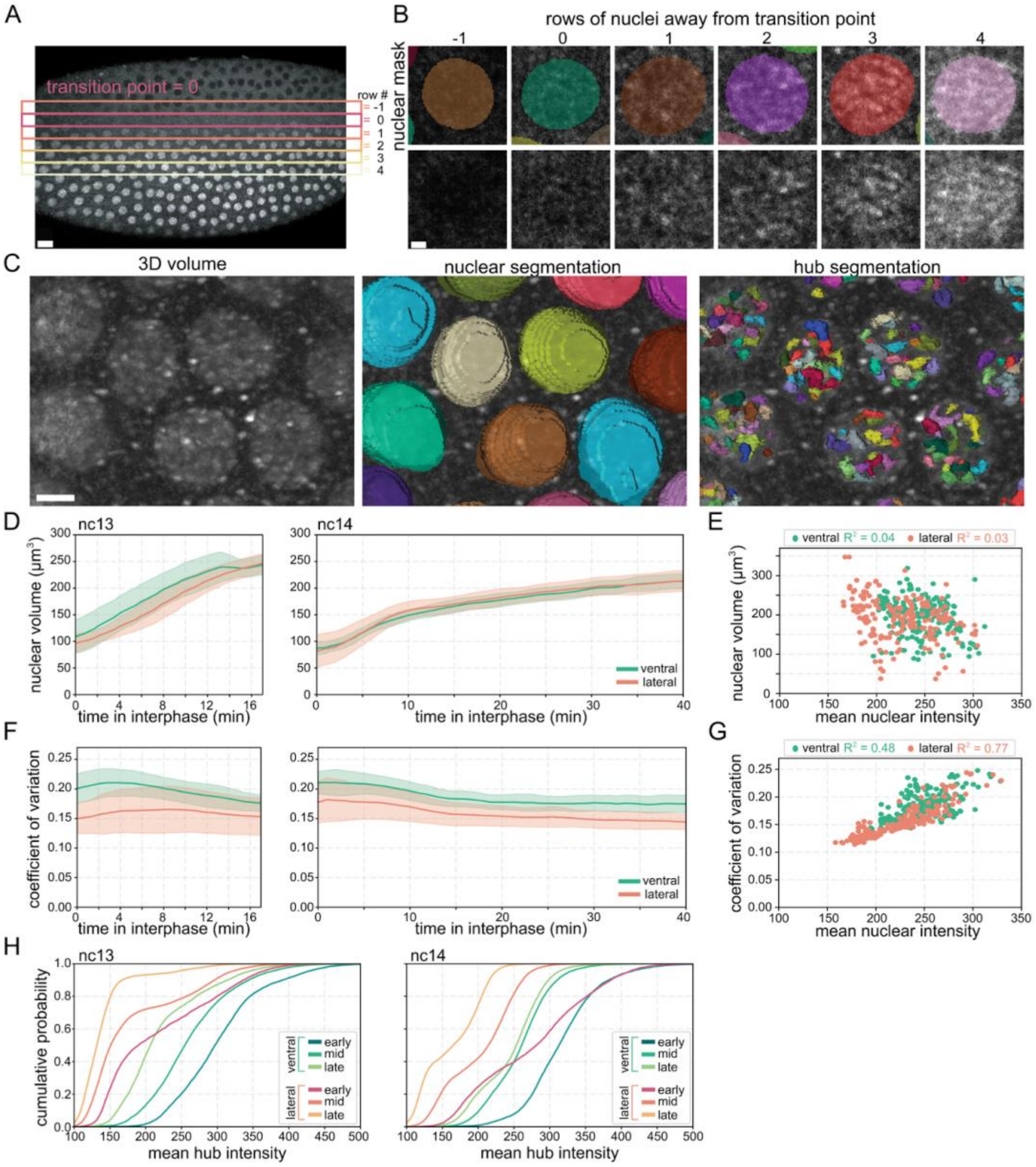


**Figure S1: Nuclear segmentation and hub detection across the dorsoventral axis.** **(A, B)** Images showing where nuclei were selected at the lateral surface within a max projection of a full field of view **(A)** and for a single slice for each nucleus. **(B)** Nuclei labelled as lateral were imaged within five nuclear rows of the transition point (row 0 of nuclei where the cytoplasm and nuclear signals are indistinguishable). Each row has a varying nuclear to cytoplasmic ratio of signal yet all have hubs within the nucleus. Scale bar in B is 15 microns and 1 micron in C. **(C)** Images showing results of nuclear and hub segmentation. Left image shows a representative 3D rendering of Dorsal-mNeonGreen in ventral nuclei. Scale bar is 2 microns. Center image shows a 3D rendering of nuclear segmentation results. Right image shows segmentation of Dorsal hubs. **(D)** Average nuclear volume in ventral (green) or lateral (orange) nuclei. Shading shows standard deviation between embryo replicates. N = 5 nuclei per embryo, 3 embryo replicates for each surface. **(E)** Scatter plot showing nuclear volume and mean nuclear intensity. $R^2$ is Pearson's correlation. **(F)** Coefficient of variation per nucleus in ventral (green) or lateral (orange) nuclei. **(G)** Scatter plot showing coefficient of variation and mean nuclear intensity. $R^2$ is Pearson's correlation. **(H)** Cumulative probability distributions of mean hub intensity for early, mid, and late nc13 (left) and nc14 (right).

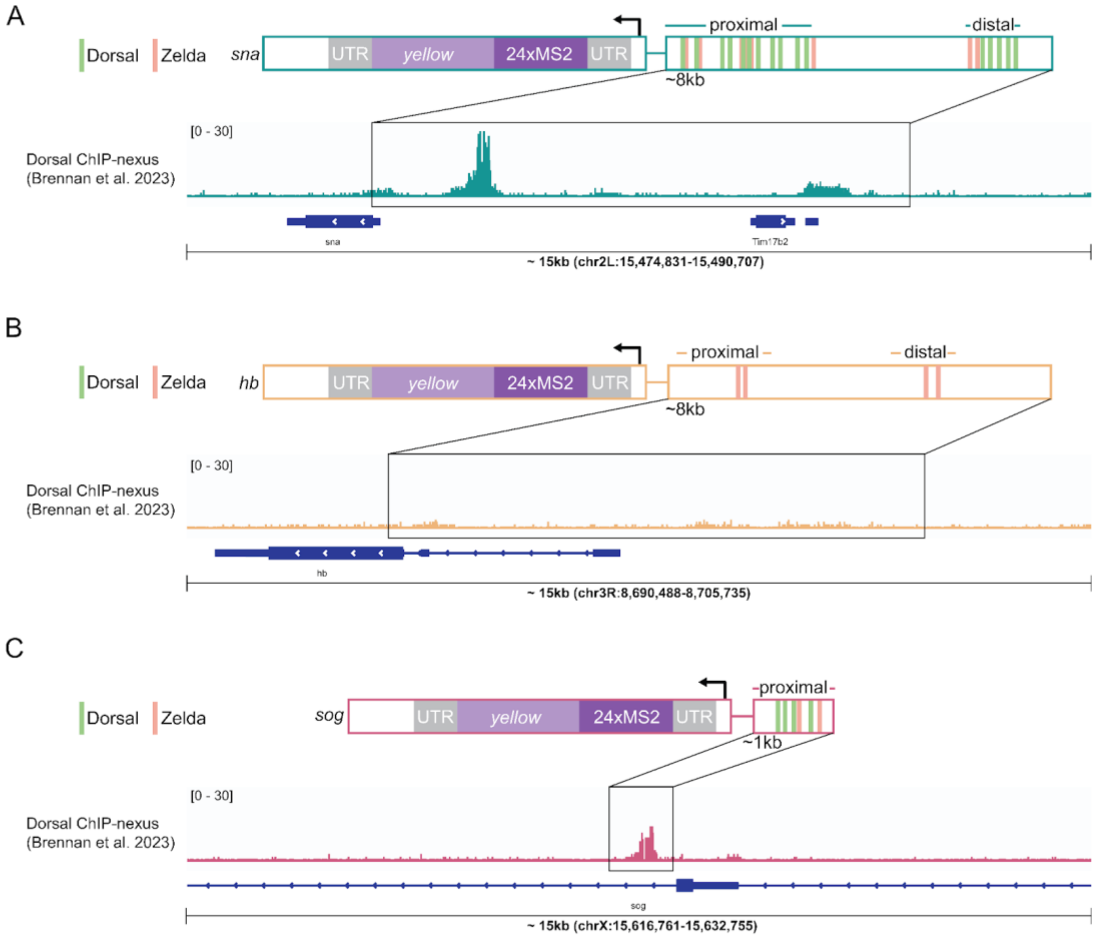


**Figure S2: Design of MS2 reporters and Dorsal occupancy in enhancer regions.** **(A)** Design of *snail-MS2 (BAC)* from (*42*) along with Dorsal occupancy (Dorsal ChIP-nexus from (*80*)) at the endogenous enhancer region corresponding to the region in the reporter. **(B)** Design of *hunchback-MS2 (BAC)* along with Dorsal occupancy at the endogenous enhancer region corresponding to the region in the reporter. **(C)** Design of *sog-MS2* from (*43*) along with Dorsal occupancy at the endogenous enhancer region corresponding to the region in the reporter

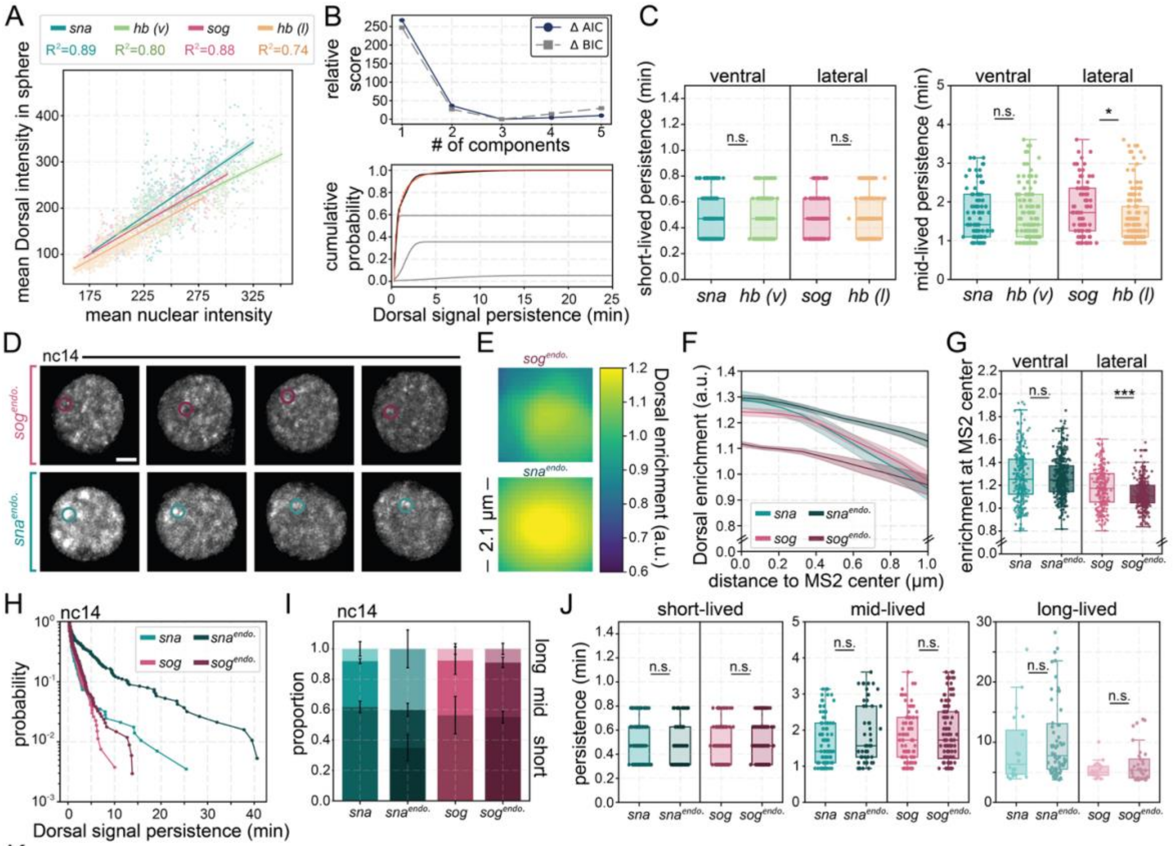


K

| | short-lived | mid-lived | long-lived |
|---|---|---|---|
| *sna* | 0.62±0.04 | 0.30±0.04 | 0.08±0.05 |
| *sna*$^{endo.}$ | 0.35±0.09 | 0.25±0.04 | 0.40±0.12 |
| *sog* | 0.56±0.12 | 0.36±0.09 | 0.08±0.03 |
| *sog*$^{endo.}$ | 0.55±0.04 | 0.36±0.03 | 0.09±0.04 |

**Figure S3. Dorsal hubs at transgenic and endogenous enhancers correlate with nuclear intensity and exhibit a range of persistence. (A)** Scatterplots show correlation between mean Dorsal intensity at each gene locus and mean nuclear intensity. $R^2$ is Pearson's Correlation. **(B)** The cumulative probability of Dorsal persistence at a locus is fit to a three-component gaussian mixture model based on minimization of Akaike Information Criterion (AIC) and Bayesian Information Criterion (BIC) (top). The fits for each component and all three together are shown (bottom). **(C)** Box plots showing the average duration of consistent hub interactions within the short- and mid-lived populations determined by a three-component gaussian-mixture model. Each persistence event is plotted. A Mann-Whitney U-test was performed to determine significance and following p-values were used: *p<0.05, **p<0.01, and ***p<0.001. **(D)** Representative maximum intensity projections of single nuclei with Dorsal-mNeonGreen in nc14. The MS2/MCP spot for each endogenous knock-in is circled in each image. The cytoplasmic Dorsal signal is masked, and contrast was adjusted independently in each row to aid visualization. Scale bars are 2 microns. **(E)** Images show average Dorsal enrichment centered at the MS2/MCP site. Images are 2.1 μm x 2.1 μm. The color bar on the bottom shows the level of Dorsal enrichment in the image (Dorsal intensity over nuclear mean intensity). **(F)** Average radial profile with standard error centered at the MS2/MCP spot shows Dorsal enrichment as a function of distance from MS2/MCP spot center. Both transgenic enhancers (Fig. 2) and endogenous knock-ins of MS2 loops for each gene are analyzed together (see Table S1 for genotypes of embryo imaged). **(G)** Boxplots showing the distribution of the Dorsal enrichment at the center of MS2/MCP site (radius = 0.0 μm). A Mann-Whitney U-test was performed to determine significance and following p-values were used: *p<0.05, **p<0.01, and ***p<0.001. **(H)** Survival plots showing the length of time any hub is consistently present within the interaction sphere for each reporter for nc14. **(I)** Using a Gaussian Mixture Model, each hub persistence event was categorized into short-, mid-, or long-lived bins. The proportion in each bin is shown with error bars designating difference among embryo replicates. **(J)** Box plots showing the average duration of consistent hub interactions within the short-, mid-, and long-lived populations determined by a three-component gaussian-mixture model. The short-lived population is bound by distinct frames acquired 9.56 seconds apart. Each persistence event from the following nuclei are plotted. N = *sna*$^{endo.}$*:* 24 nuclei in 3 embryos and *sog*$^{endo.}$*:* 24 nuclei in 3 embryos. A Mann-Whitney U-test was performed to determine significance. **(K)** The proportion of short-, mid-, and long-lived hub persistence events are listed with their mean and standard deviation in the table.

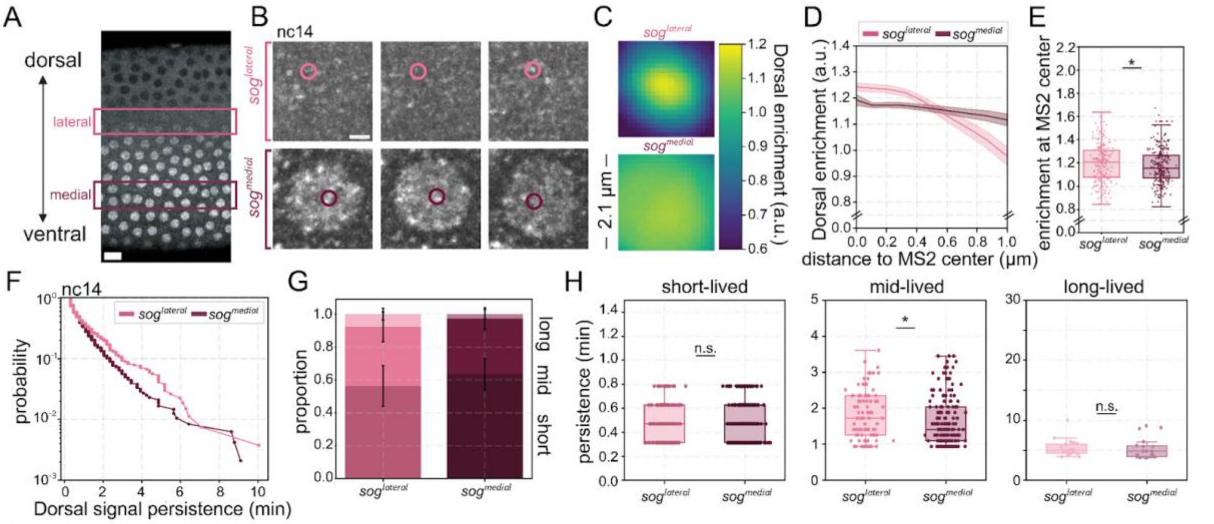


I

| | short-lived | mid-lived | long-lived |
|---|---|---|---|
| $sog^{lateral}$ | 0.56±0.12 | 0.36±0.09 | 0.08±0.03 |
| $sog^{medial}$ | 0.63±0.10 | 0.34±0.07 | 0.03±0.03 |

**Figure S4: Differences in apparent Dorsal hub properties at *sog* at different Dorsal concentrations is due to nuclear background concentration.** **(A)** Image of Dorsal-mNeonGreen where the 'lateral' and 'medial' *sog* nuclei were imaged (see Table S1 for genotypes of embryo imaged). **(B)** Representative maximum intensity projections of single nuclei with Dorsal-mNeonGreen in nc14. The *sog*-MS2 reporter location is circled in each image. The scale bar is 2 microns. **(C)** Images show average Dorsal enrichment centered at the MS2/MCP site. Images are 2.1 µm x 2.1 µm. The color bar on the bottom shows the level of Dorsal enrichment in the image (Dorsal intensity over nuclear mean intensity). **(D)** Average radial profile with standard error centered at the MS2/MCP spot shows Dorsal enrichment as a function of distance from MS2/MCP spot center. **(E)** Boxplots showing the distribution of the Dorsal enrichment at the center of MS2/MCP site (radius = 0.0 µm). A Mann-Whitney U-test was performed to determine significance and following p-values were used: *p<0.05, **p<0.01, and ***p<0.001. **(F)** Survival plots showing the length of time any hub is consistently present within the interaction sphere for each reporter for nc14. **(G)** Using a Gaussian Mixture Model, each hub persistence event was categorized into short-, mid-, or long-lived bins. The proportion in each bin is shown with error bars designating standard deviation among embryo replicates. **(H)** Box plots showing the average duration of consistent hub interactions within the short-, mid-, and long-lived populations determined by a three-component gaussian-mixture model. The short-lived population is bound by distinct frames acquired 9.56 seconds apart. Each persistence event from the following nuclei are plotted. N = $sog^{lateral}$: 18 nuclei in 3 embryos and $sog^{medial}$: 24 nuclei in 3 embryos. A Mann-Whitney U-test was performed to determine significance and following p-values were used: *p<0.05, **p<0.01, and ***p<0.001. **(I)** The proportion of short-, mid-, and long-lived hub persistence events are listed with their mean and standard deviation in the table.

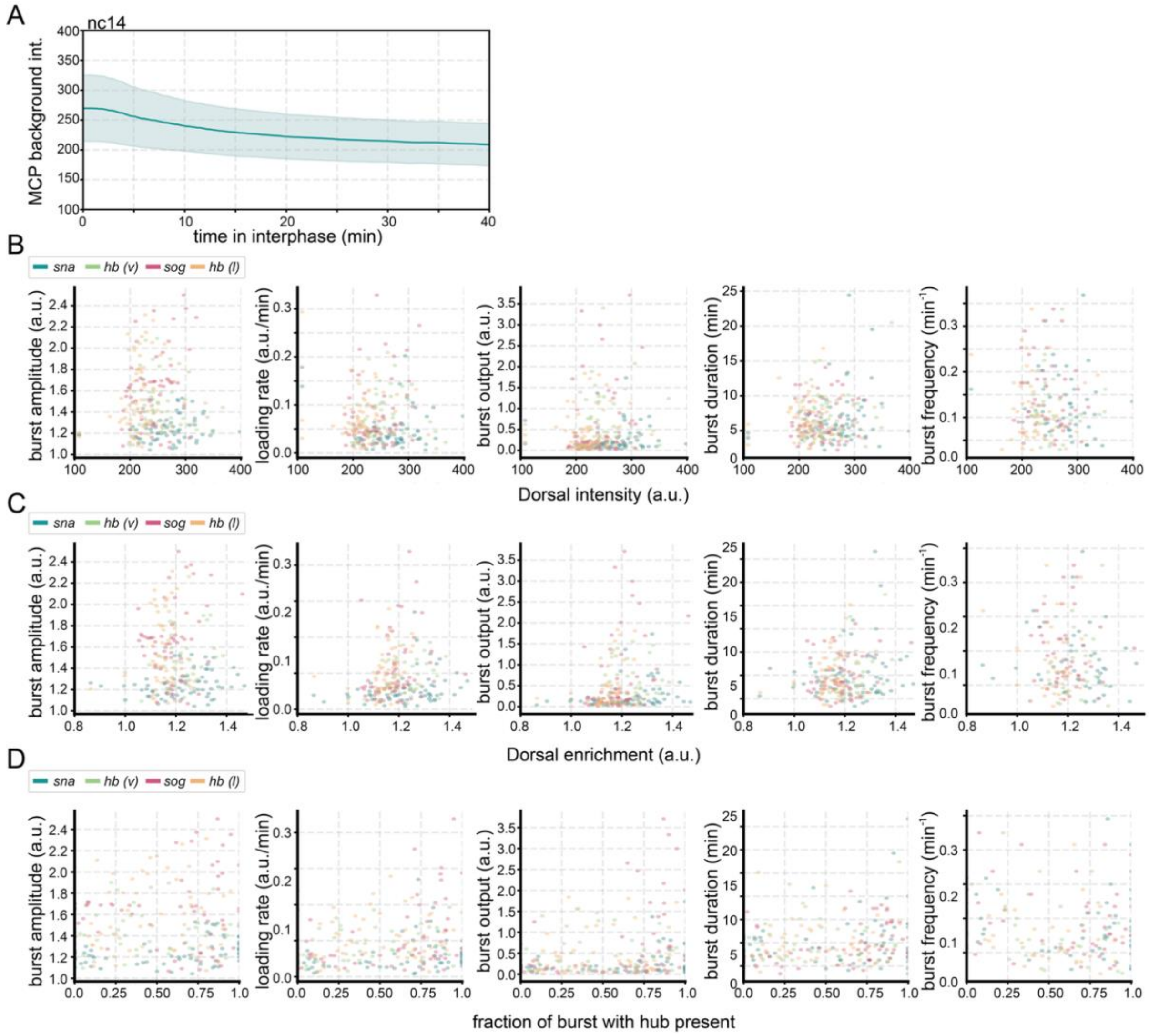


**Figure S5: Hub properties and transcription burst kinetics show minimal correlations.** **(A)** Average MCP background intensity within nuclei. MCP intensity decreases ~77% during nc14 due to photobleaching. **(B-D)** Scatter plots showing burst metrics (amplitude, loading rate, output, duration, frequency) with the mean Dorsal intensity **(B)**, Dorsal enrichment **(C)**, or fraction of burst with a hub present **(D)** before the burst (left), the mean Dorsal enrichment before the burst (center), or the fraction of time during a burst when a hub is persisting near the MS2/MCP site (right). Marker colors correspond to each reporter analyzed. These data points were used in the heat maps reflecting Spearman's rank correlation in Figure 3D-F.

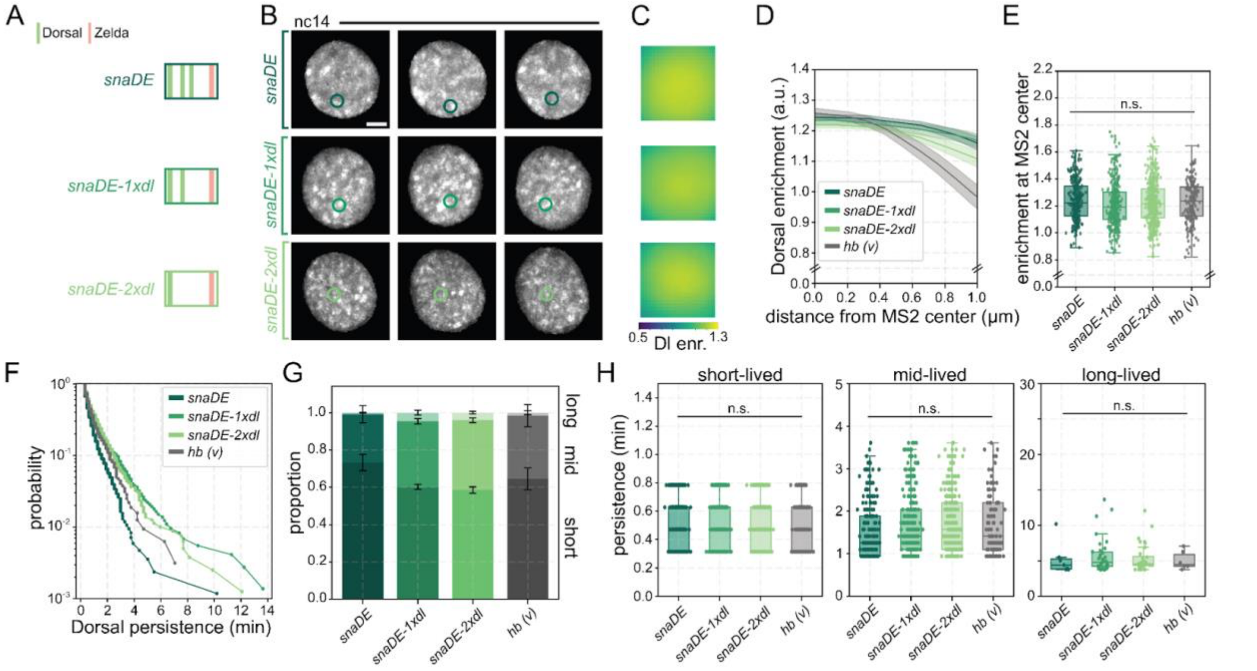


**Figure S6: Hub enrichment and signal persistence detection limits.** **(A)** Schematic depicting enhancers with Dorsal and Zelda binding sites for each *snail* reporter with the minimal distal enhancer of *snail* (*snaDE*), minus one Dorsal binding site (*snaDE-1xdl*), or minus two Dorsal binding site (*snaDE-2xdl*). Each enhancer was positioned just upstream of its promoter, 10 MS2 stem loops, and the *yellow* coding sequence (see Table S1 for exact genotypes of embryo imaged). Each of these *snail* reporters were imaged along the ventral surface in nc14. **(B)** Representative maximum intensity projections of single nuclei with Dorsal-mNeonGreen in nc14. The MS2/MCP site is circled in each image. The cytoplasmic Dorsal signal is masked, and contrast was adjusted independently in each row to aid visualization. Scale bars are 2 microns. **(C)** Images show average Dorsal enrichment centered at the MS2/MCP site. Images are 2.1 μm x 2.1 μm. The color bar on the bottom shows the level of Dorsal enrichment in the image (Dorsal intensity over nuclear mean intensity). **(D)** Average radial profile with standard error centered at the MS2/MCP spot shows Dorsal enrichment as a function of distance from MS2/MCP spot center. **(E)** Boxplots showing the distribution of the Dorsal enrichment at the center of MS2/MCP site (radius = 0.0 μm). A Mann-Whitney U-test was performed to determine significance. **(F)** Survival plots showing the length of time any hub is consistently present within the interaction sphere for each reporter for nc14. **(G)** After applying a Gaussian Mixture Model, each hub persistence event was categorized into short-, mid-, or long-lived bins. The proportion in each bin is shown with error bars designating standard deviation among embryo replicates. **(H)** Box plots showing the average duration of consistent hub interactions within the short-, mid-, and long-lived populations determined by a three-component gaussian-mixture model. The short-lived population is bound by distinct frames acquired 9.56 seconds apart. Each persistence event from the following nuclei are plotted. N = *snaDE:* 28 nuclei in 3 embryos; *snaDE-1xdl:* 33 nuclei in 3 embryos; *snaDE-2xdl:* 44 nuclei in 4 embryos; and *hb (v):* 18 nuclei in 4 embryos. A Mann-Whitney U-test was performed to determine significance and following p-values were used: *p<0.05, **p<0.01, and ***p<0.001

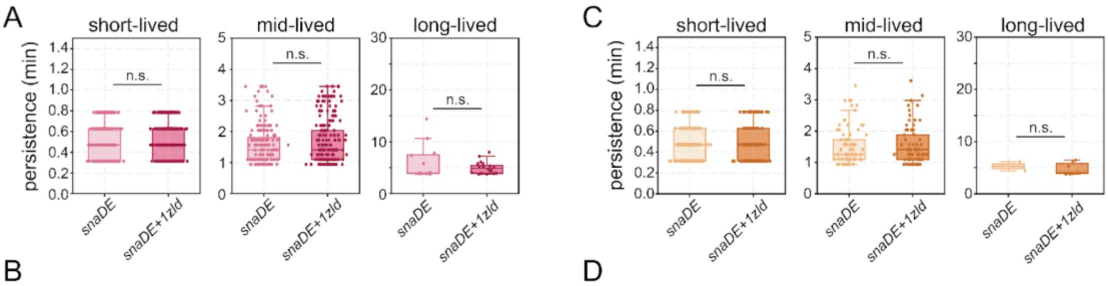


B

| Dorsal | short-lived | mid-lived | long-lived |
|---|---|---|---|
| *snaDE* | 0.66±0.04 | 0.32±0.04 | 0.02±0.01 |
| *snaDE+1zld* | 0.64±0.10 | 0.33±0.08 | 0.03±0.03 |

D

| Zelda | short-lived | mid-lived | long-lived |
|---|---|---|---|
| *snaDE* | 0.72±0.10 | 0.28±0.10 | 0.01±0.01 |
| *snaDE+1zld* | 0.73±0.03 | 0.25±0.04 | 0.02±0.01 |

**Figure S7: Dorsal and Zelda persistence are moderately impacted from a single Zelda binding site addition.** **(A, C)** Box plots showing the average duration of consistent Dorsal **(A)** or Zelda **(C)** hub interactions within the short-, mid-, and long-lived populations determined by a three-component gaussian-mixture model. The short-lived population is bound by distinct frames acquired 9.56 seconds apart. Each persistence event from all nuclei are plotted. A Mann-Whitney U-test was performed to determine significance and following p-values were used: *$p<0.05$, **$p<0.01$, and ***$p<0.001$. **(B, D)** The proportion of short-, mid-, and long-lived Dorsal **(B)** or Zelda **(D)** hub persistence events are listed with their means and standard deviations in the table.

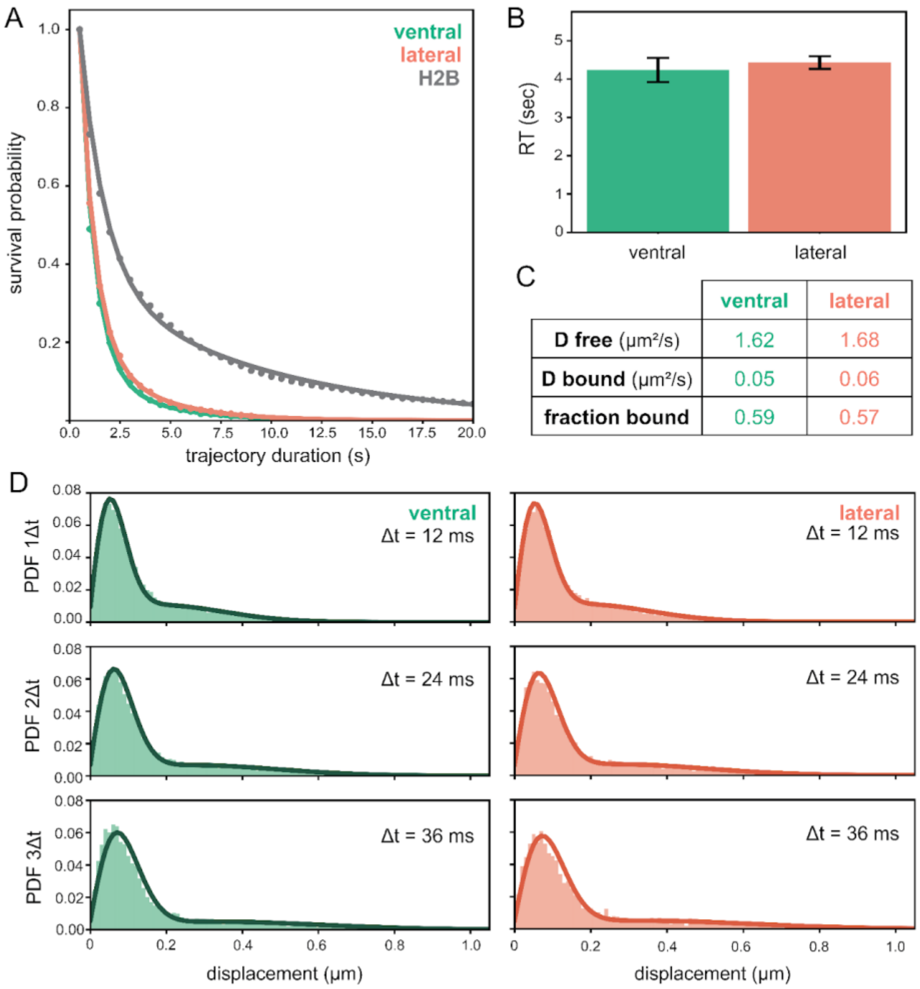


**Figure S8: Single molecule tracking (SMT) of Dorsal shows minimal differences on the ventral and lateral regions. (A)** Survival probabilities of single-molecule trajectories at 500 msec exposure times for at least three embryos for Dorsal-mEos4a ventrally positioned (N = 978 trajectories, 5 unique embryos), Dorsal-mEos4a laterally positioned (N = 1214 unique trajectories, 4 unique embryos), and H2B-mEos4a (N = 3098 trajectories, 3 unique embryos)**. (B)** Doral residence times after photobleaching correction for ventral (4.24 ± 0.32 sec) and lateral (4.43 ± 0.17 sec) positions. **(C)** Spot-On diffusion coefficient estimates for free and bound populations. **(D)** Probability density functions (PDFs) of the trajectory displacements of ventral (N = 10968 trajectories, 4 unique embryos) and lateral (N = 5824 trajectories, 11 unique embryos) Dorsal-mEos4a across three time intervals.

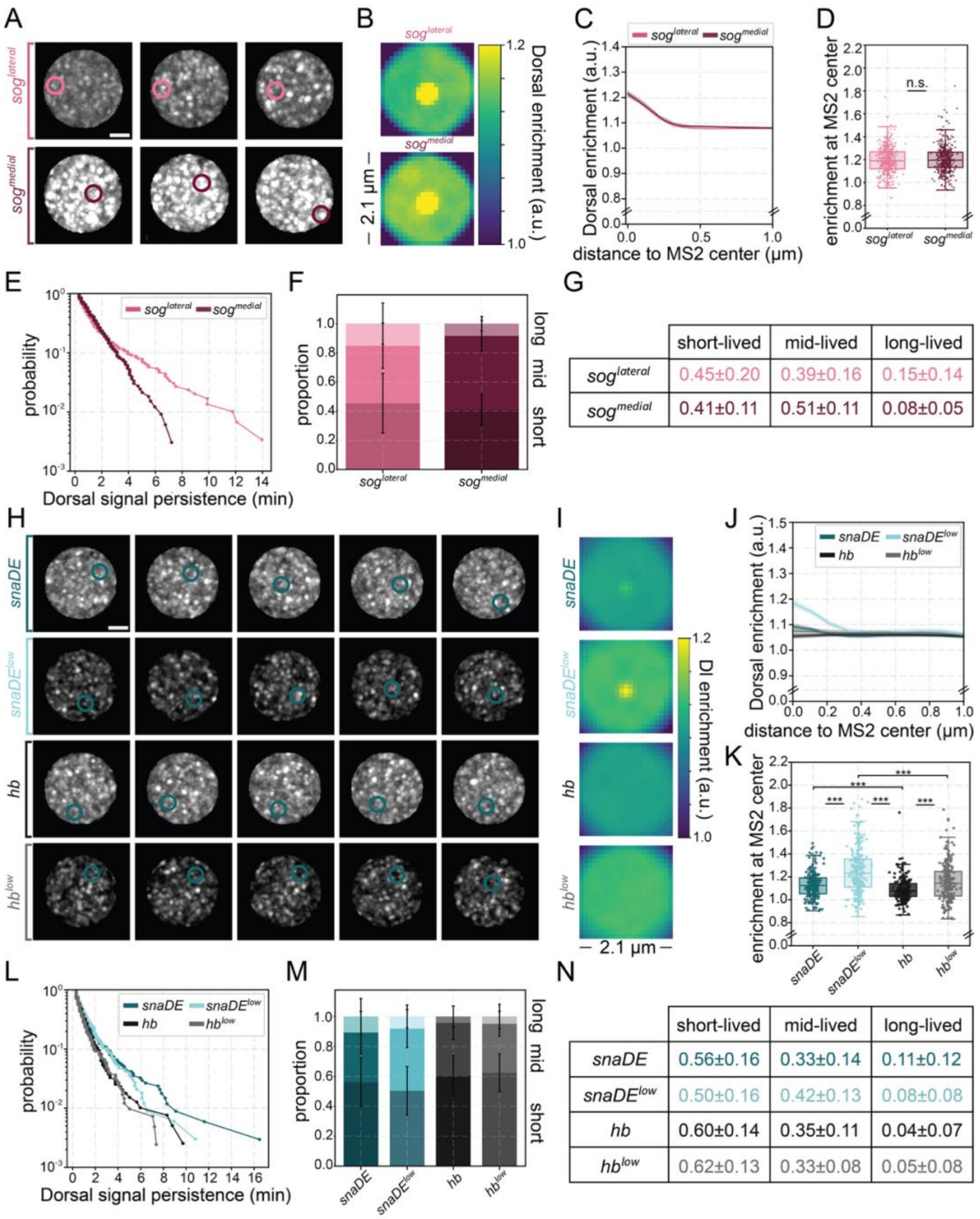


**Figure S9: Computational modeling shows the effect of nuclear concentration and imaging conditions on hub properties.** **(A)** Representative maximum intensity projections of simulated single nuclei with Dorsal signal in grey. The binding site cluster is circled in each image. Contrast is scaled equally for all conditions. Scale bars are 2 microns. **(B)** Images show simulated average Dorsal enrichment centered at the binding site clusters. Images are 2.1 µm x 2.1 µm. The color bar on the bottom shows the level of Dorsal enrichment in the image (Dorsal intensity over nuclear mean intensity). **(C)** Average radial profile with standard error centered at the binding site cluster shows Dorsal enrichment as a function of distance from spot center. **(D)** Boxplots showing the distribution of the Dorsal enrichment at the center of the binding site cluster (radius = 0.0 µm). A Mann-Whitney U-test was performed to determine significance and following p-values were used: *p<0.05, **p<0.01, and ***p<0.001. **(E)** Survival plots showing the length of time any hub is consistently present within the interaction sphere for each reporter. **(F)** The proportion of long-, mid-, and short-lived persistent hub formation in each bin is shown with error bars designating standard deviation among embryo replicates. **(G)** The proportion of short-, mid-, and long-lived hub persistence events are listed with their mean and standard deviation in the table. **(H)** Representative maximum intensity projections of simulated single nuclei with Dorsal signal in grey. The binding site cluster is circled in each image. Contrast is scaled equally for all conditions. Scale bars are 2 microns. Simulations show *snaDE* or *hb* visualized with regular imaging conditions or with lowered background and noise. **(I)** Images show simulated average Dorsal enrichment centered at the binding site clusters. Images are 2.1 µm x 2.1 µm. The color bar on the bottom shows the level of Dorsal enrichment in the image (Dorsal intensity over nuclear mean intensity). **(J)** Average radial profile with standard error centered at the binding site cluster shows Dorsal enrichment as a function of distance from spot center. **(K)** Boxplots showing the distribution of the Dorsal enrichment at the center of the binding site cluster (radius = 0.0 µm). A Mann-Whitney U-test was performed to determine significance and following p-values were used: *p<0.05, **p<0.01, and ***p<0.001. **(L)** Survival plots showing the length of time any hub is consistently present within the interaction sphere for each reporter. **(M)** The proportion of long-, mid-, and short-lived persistent hub formation in each bin is shown with error bars designating standard deviation among embryo replicates. **(N)** The proportion of short-, mid-, and long-lived hub persistence events are listed with their mean and standard deviation in the table.

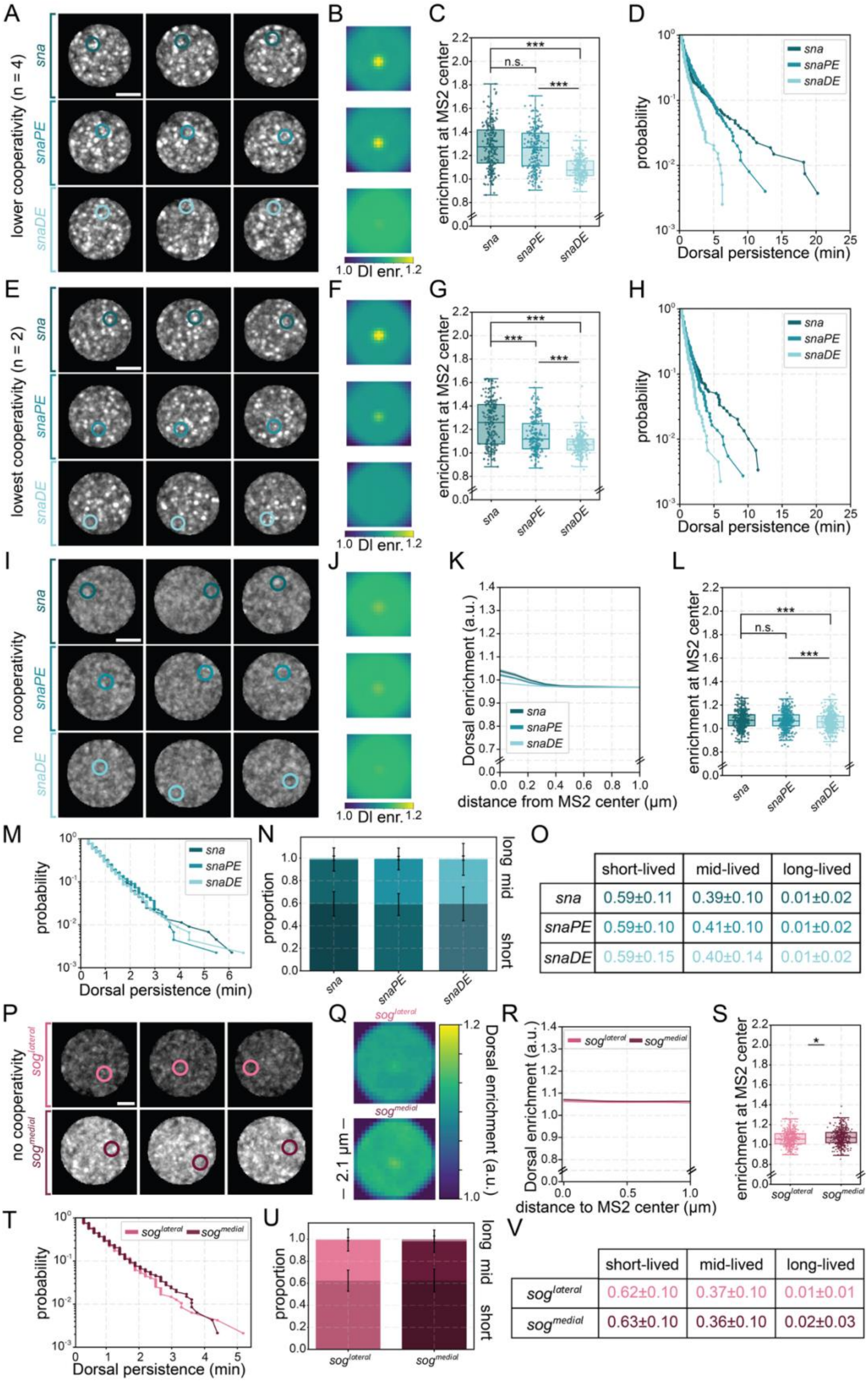


**Figure S10: Computational modeling without cooperativity fails to recapitulate Dorsal hub properties.** **(A, E, I P)** Representative maximum intensity projections of simulated single nuclei with cooperative binding (Hill-like = 2 **(A)**, 4 **(E)**) or without cooperative binding included in computation **(I, P)** (Dorsal signal in grey). The binding site cluster is circled in each image. Contrast is scaled equally for all conditions. Scale bars are 2 microns. **(B, F, J, Q)** Images show simulated average Dorsal enrichment centered at the binding site clusters. Images are 2.1 μm x 2.1 μm. The color bar on the bottom shows the level of Dorsal enrichment in the image (Dorsal intensity over nuclear mean intensity). **(C, G, L, S)** Boxplots showing the distribution of the Dorsal enrichment at the center of the binding site cluster (radius = 0.0 μm). A Mann-Whitney U-test was performed to determine significance and following p-values were used: *p<0.05, **p<0.01, and ***p<0.001. **(D, H, M, T)** Survival plots showing the length of time any hub is consistently present within the interaction sphere for each reporter. **(K, R)** Average radial profile with standard error centered at the binding site cluster shows Dorsal enrichment as a function of distance from spot center. **(N, U)** The proportion of long-, mid-, and short-lived persistent hub formation in each bin is shown with error bars designating standard deviation among embryo replicates. **(O, V)** The proportion of short-, mid-, and long-lived hub persistence events are listed with their mean and standard deviation in the table

### Table S1: Details of *Drosophila* lines used.

| **Shorthand** | **Genotype*** | **Source** | **Appears in Figures** |
|---|---|---|---|
| Dorsal-mNeonGreen | yw; Dorsal-mNeonGreen/CyO; | (*41*) | 1, 2, 3, 4, S1, S3, S4, S5, S6, S7, S8 |
| Dorsal-mEos4a | yw; Dorsal-mEos4a/CyO; | (*41*) | 6, S8 |
| mNeonGreen-Zelda | mNeonGreen-Zelda;; | (*52*) | 5, S7 |
| MCP-mCherry | nos>MCP-mCherry (III) | (*54*) | 2, 3, 4, S3, S4, S5, S6, S7, S8 |
| MCP-mCherry | enosx2>MCP-mCherry (II) | (*41*) | 5, S7 |
| *sna* | sna-24xMS2 (BAC) | (*42*) | 2, 3, S3, S5 |
| *hb* | hb-24xMS2 (BAC) | (*42*) | 2, 3, 4, S3, S5 |
| *sog*, $sog^{lateral}$, $sog^{medial}$ | sog-24xMS2 (BAC) | (*43*) | 2, 3, S3, S4, S5 |
| $sna^{endo.}$ | sna-24xMS2 (endogenous 3') | (*44*) | S3 |
| $sog^{endo.}$ | sog-24xMS2 (endogenous 5') | Gift from Bomyi Lim | S3 |
| *snaPE* | snaPE > snaPr > 24xMS2-yellow | Gift from Bomyi Lim | 4 |
| *snaDE* | snaDE > snaPr > 24xMS2 - yellow | (*39*) | 4, 5, S7 |
| *snaDE* | snaDE > snaPr > 10xMS2 - yellow | (*49*) | S6 |
| *snaDE - 1xdl* | snaDE del dl1 > snaPr > 10xMS2 - yellow | (*49*) | S6 |
| *snaDE - 2xdl* | snaDE del dl1/2 > snaPr > 10xMS2 - yellow | (*49*) | S6 |
| *snaDE + 1 zld* | snaDE + 1 Zld 5' > snaPr > 24xMS2 - yellow | (*39*) | 5, S7 |

* All non-endogenous MS2 reporters inserted into VK33 on Chromosome III

### Table S2: Simulation parameters

| Category | Parameter | Value |
|---|---|---|
| General Parameters | Dorsal concentration from (*54*) | Ventral = 500 nM |
| | | Medial = 406 nM |
| | | Lateral = 219 nM |
| | Nucleus diameter | 5 µm |
| | Molecule diameter | 15 nm |
| | Simulation time step | 10 ms |
| | Display time step | 10 s |
| Binding sites | Number (post-replication) from (*50*) | 7000 |
| | Endogenous sites per cluster | 2 (60%), 10 (25%), 20 (10%), 30 (4%), 40 (1%) |
| | Endogenous site cluster diameter | 3 px = 324 nm |
| | Diffusion coefficient | 0.05 ± 0.03 $\mu m^2/s$ (0.001 µm jiggle) |
| | Hurst exponent | 0.25 |
| | Synthetic sites per cluster (post-replication) | *snail* = 32 (in a 1, 3 or 5 px diameter) |
| | | *snailPE* = 22 (in a 1, 2 or 4 px diameter) |
| | | *snailDE* = 6 (in a 1, 2 or 3 px diameter) |
| | | *sog* = 8 (in a 1, 2 or 3 px diameter) |
| Three-state diffusion model | Initialization | 50% free, 50% non-specifically bound |
| | Free diffusion coefficient | 1.7 ± 0.45 $\mu m^2/s$ |
| | Switching rate (free to non-specific binding) | 1 Hz |
| | Non-specific residence time | 0.5 s |
| | Binding probability, $k_{on}$ | 0.005 |
| | Capture radius | 50 nm |
| Binding site strength distribution | Base residence time, $1/k_{off}$ | 5 ± 1 s |
| | Endogenous site residence time (multipliers of base residence time) | 0.06x (50%), 0.2x (27%), 0.5x (15%), 1x (6%), 2.4x (2%) |
| | Synthetic site residence time (multipliers of base residence time) | *snail*/*snailPE*/*snailDE* = 0.2x |
| | | *hunchback* = 0.000001x |
| | | *sog* = 0.5x |
| | Hill-like coefficient, $n$ | 2 and 4 (lowered cooperativity), or 100 (with cooperativity) |

### Supplemental Movie Captions.

**Movie S1. Dorsal hub formation in nc13 and 14.**
Movies showing Dorsal-mNeonGreen (gray) in nuclei along the lateral (left) or ventral (right) surface of a *Drosophila* embryo in nc13 and nc14. Each movie is a 3D rendering with equivalent contrast limits.

**Movie S2. Dorsal hubs at the lateral transition point.**
Movie showing Dorsal-mNeonGreen (gray) in 3D rendered nuclei along the lateral transition point of a *Drosophila* embryo in nc13 and nc14.

**Movie S3. Dorsal hub segmentation example.**
Example of hub segmentation (colored labels) within ventral nuclei of Dorsal-mNeonGreen (gray). Movie starts on a single z-slice then moves through the whole nucleus in z.

**Movie S4. Dorsal hubs at target and non-target genes in nc13.**
Example 3D rendering of nuclei through time showing Dorsal-mNeonGreen (gray) at MS2 spots (circled) for *sna, hb (v), sog,* and *hb (l)* in nc13. For visualization purposes, cytoplasm has been removed and set to zero, and contrast limits are individually adjusted for ventral or lateral nuclei.

**Movie S5. Dorsal hubs at target and non-target genes in nc14.**
Example 3D rendering of nuclei through time showing Dorsal-mNeonGreen (gray) at MS2 spots (circled) for *sna, hb (v), sog,* and *hb (l)* in nc14. For visualization purposes, cytoplasm has been removed and set to zero, and contrast limits are individually adjusted for ventral or lateral nuclei.

**Movie S6. Example of MS2/MCP tracking at *sog*.**
Example of 3D MS2 tracking in time within a single nucleus. Gray channel shows MCP-mCherry distribution in the nucleus. For visualization purposes, cytoplasm has been removed and set to zero. A pink circle marks the spot of nascent transcription of *sog*-MS2. The left shows a top-down (X-Y) view while the right shows a side (Y-Z) view.

**Movie S7. Dorsal hubs at endogenous target genes.**
Example 3D rendering of nuclei through time showing Dorsal-mNeonGreen (gray) at MS2 spots (circled) for *snail (endogenous)* and *sog (endogenous)* in nc14. For visualization purposes, cytoplasm has been removed and set to zero, and contrast limits are individually adjusted for ventral or lateral nuclei.

**Movie S8. Dorsal hubs at *sog* at different concentrations.**
Example 3D rendering of nuclei through time showing Dorsal-mNeonGreen (gray) at MS2 spots (circled) for *sog (lateral)* and *sog (medial)* in nc14. Contrast limits are equivalent across conditions.

**Movie S9. Dorsal hubs at various *snail* enhancers.**
Example 3D rendering of nuclei through time showing Dorsal-mNeonGreen (gray) at MS2 spots (circled) for *sna, snaPE,* and *snaDE* in nc14. For visualization purposes, cytoplasm has been removed and set to zero. Contrast limits are equivalent across conditions.

**Movie S10. Dorsal hubs when binding sites are removed.**
Example 3D rendering of nuclei through time showing Dorsal-mNeonGreen (gray) at MS2 spots (circled) for *snaDE, snaDE-1xdl,* and *snaDE-2xdl* in nc14. For visualization purposes, cytoplasm has been removed and set to zero. Contrast limits are equivalent across conditions.

**Movie S11. Dorsal hubs upon the addition of a Zelda binding site.**
Example 3D rendering of nuclei through time showing Dorsal-mNeonGreen (gray) at MS2 spots (circled) for *snaDE* and *snaDE+1zld* in nc14. For visualization purposes, cytoplasm has been removed and set to zero. Contrast limits are equivalent across conditions.

**Movie S12. Zelda hubs upon the addition of a Zelda binding site.**
Example 3D rendering of nuclei through time showing mNeonGreen-Zelda (gray) at MS2 spots (circled) for *snaDE* and *snaDE+1zld* in nc14. For visualization purposes, cytoplasm has been removed and set to zero. Contrast limits are equivalent across conditions.
